# A hidden Markov model approach for simultaneously estimating local ancestry and admixture time using next generation sequence data in samples of arbitrary ploidy

**DOI:** 10.1101/064238

**Authors:** Russell Corbett-Detig, Rasmus Nielsen

**Affiliations:** Department of Biomolecular Engineering, UC Santa Cruz, Santa Cruz, CA; Department of Integrative Biology, UC Berkeley, Berkeley, CA; The Natural History Museum of Denmark, University of Copenhagen, Denmark.

**Keywords:** Local Ancestry Inference, Admixture, Hidden Markov Model, *Drosophila melanogaster*, Pool-seq

## Abstract

Admixture—the mixing of genomes from divergent populations—is increasingly appreciated as a central process in evolution. To characterize and quantify patterns of admixture across the genome, a number of methods have been developed for local ancestry inference. However, existing approaches have a number of shortcomings. First, all local ancestry inference methods require some prior assumption about the expected ancestry tract lengths. Second, existing methods generally require genotypes, which is not feasible to obtain for many next-generation sequencing projects. Third, many methods assume samples are diploid, however a wide variety of sequencing applications will fail to meet this assumption. To address these issues, we introduce a novel hidden Markov model for estimating local ancestry that models the read pileup data, rather than genotypes, is generalized to arbitrary ploidy, and can estimate the time since admixture during local ancestry inference. We demonstrate that our method can simultaneously estimate the time since admixture and local ancestry with good accuracy, and that it performs well on samples of high ploidy—*i.e.* 100 or more chromosomes. As this method is very general, we expect it will be useful for local ancestry inference in a wider variety of populations than what previously has been possible. We then applied our method to pooled sequencing data derived from populations of *Drosophila melanogaster* on an ancestry cline on the east coast of North America. We find that regions of local recombination rates are negatively correlated with the proportion of African ancestry, suggesting that selection against foreign ancestry is the least efficient in low recombination regions. Finally we show that clinal outlier loci are enriched for genes associated with gene regulatory functions, consistent with a role of regulatory evolution in ecological adaptation of admixed *D. melanogaster* populations. Our results illustrate the potential of local ancestry inference for elucidating fundamental evolutionary processes.

**Author Summary:** When divergent populations hybridize, their offspring obtain portions of their genomes from each parent population. Although the average ancestry proportion in each descendant is equal to the proportion of ancestors from each of the ancestral populations, the contribution of each ancestry type is variable across the genome. Estimating local ancestry within admixed individuals is a fundamental goal for evolutionary genetics, and here we develop a method for doing this that circumvents many of the problems associated with existing methods. Briefly, our method can use short read data, rather than genotypes and can be applied to samples with any number of chromosomes. Furthermore, our method simultaneously estimates local ancestry and the number of generations since admixture—the time that the two ancestral populations first encountered each other. Finally, in applying our method to data from an admixture zone between ancestral populations of *Drosophila melanogaster*, we find many lines of evidence consistent with natural selection operating to against the introduction of foreign ancestry into populations of one predominant ancestry type. Because of the generality of this method, we expect that it will be useful for a wide variety of existing and ongoing research projects.

## Introduction

Characterizing the biological consequences of admixture—the mixing of genomes from divergent ancestral populations—is a fundamental and important challenge in evolutionary genetics. Admixture has been reported in a variety of natural populations of animals [1,2], plants [3–5] and humans [6,7], and theoretical and empirical evidence suggests that admixture may affect a diverse suite of evolutionary processes. Individuals’ ancestry can affect disease susceptibility in admixed populations, and inferring and correcting for sample population ancestries is a common practice in human genome wide association studies [8–10]. More generally, admixture has the potential to influence patterns of genetic variation within populations [11,12], to introduce novel adaptive [13,14] and deleterious variants [7,15,16], as well as to disrupt epistatic gene networks [17,18]. Therefore, developing a comprehensive understanding of the extent of admixture in natural populations and resulting mosaic genome structures is essential to furthering our understanding of a variety of evolutionary processes.

Estimating genome-wide ancestry proportions has become a common practice in population genetic inference. For example, the program STRUCTURE [19], originally released in 2000, uses a Bayesian framework to model the ancestry proportions of individuals derived from any number of source populations based on genotype data at a set of unlinked genetic markers. More recently, this model for ancestry proportion estimation has been extended to cases where individual genotypes are not known, but can be studied probabilistically using low-coverage sequencing short read sequencing data [20], which is an important step towards accommodating modern sequencing practices. Additionally, Bergland *et. al*. [21] developed a method for estimating ancestry proportions in pooled population samples of relatively high ploidy (*i.e.* 40–250 distinct chromosomes) from short read sequencing data. In general, it is straightforward to estimate genome-wide ancestry proportions using a number of sequencing strategies and applications.

It is substantially more challenging to accurately estimate local ancestry (LA) at markers distributed along the genome of a sample. Nonetheless, analyses of LA have the potential to yield more nuanced insights into our understanding of the evolutionary processes affecting ancestry proportions across the genome. One of the first LA inference (LAI) methods was an extension of the STRUCTURE [19] framework that modeled the correlation in ancestry among markers due to linkage. Because the ancestry at each locus is not observed, Falush *et al.* [22] suggested that a hidden Markov model (HMM) is a straightforward means of inferring the ancestry states at each site in the genome (which are unobserved) based on observed genotype data distributed along a chromosome. Most subsequent LAI methods have also used an HMM framework, and the majority are geared towards estimating LA in admixed human populations (*e.g.* [23,24]). Consequently, most existing LAI methods are limited to diploid genomes with high quality genotype calls. Furthermore, many methods require phased reference panels [24,25], and require the user to provide an estimate of, or make implicit assumptions about, the number of generations since the initial admixture event [2,23–25]. This is straightforward with human population genomic samples, where abundant high quality genotyped samples are available and for which well-documented demographic histories are sometimes known. However for most other species, demographic histories are less well characterized, and assumptions about admixture times may bias the result of LAI methods.

A number of approaches exist to estimate the time since admixture based on well characterized ancestry tract length distributions [26–29] but in general, these parameters are unknown prior to LAI. Conversely, another class of methods can be used to estimate the time of admixture based on the decay of linkage disequilibrium without performing LAI [30–32]; however as with LAI procedure above, these approaches are also limited to diploid genotype data. We may therefore expect to improve LAI by simultaneously estimating LA and demographic parameters (*e.g.* admixture time). Furthermore, in the majority of sequencing applications, relatively low individual sequencing coverage is often used to probabilistically estimate individual and population allele frequencies (*e.g.* [33]) but these data are often not sufficient to determine high confidence genotypes that are required for existing LAI applications. Hence, there is a clear need for a general LAI method that can accommodate genotype uncertainty and requires less advanced knowledge of admixed populations’ demographic histories.

Here, we introduce a framework for simultaneously estimating LA using short read pileup data and the time of admixture within a population. Briefly, as with many previously proposed LAI methods, we model ancestry across the genome of a sample as a HMM. We estimate LA by explicitly modeling read counts as a function of sample allele frequencies within an admixed population. Our method is generalized to accommodate arbitrary sample ploidies, and is therefore applicable to haploid (or inbred), diploid, tetraploid, as well as pooled sequencing applications. We show that this approach accurately infers the time since admixture when data are simulated under the assumed model. Furthermore, our method yields accurate LA estimates for simulated datasets, including samples of high sample ploidy and including evolutionary scenarios that violate the assumptions of the neutral demographic model. In comparisons between ours and an existing LAI method, WINPOP [34], we find that our approach offers a significant improvement and is accurate over longer time scales. Furthermore, we demonstrate, using a published dataset, that even state-of-the-art LAI methods can be significantly impacted by assumptions about the time since admixture, and that our method provides a solution to this problem.

Finally, we apply this method to a *Drosophila melanogaster* ancestry cline on the east coast of North America. This species originated in sub-Saharan Africa, and approximately 10,000—15,000 years ago a subpopulation expanded out of the ancestral range. During this expansion, the derived subpopulation experienced a population bottleneck that resulted in decreased nucleotide polymorphism, extended linkage disequilibrium within the derived population and substantial genetic differentiation between ancestral and derived populations [2,35–39]. Hereafter, the ancestral population will be referred to as “African” and the derived population as “Cosmopolitan”. Following this bottleneck, descendant populations of African and Cosmopolitan *D. melanogaster* have admixed in numerous geographic regions [2,11,21]. Of particular relevance to this work, North America was colonized recently by a population descendent from African individuals from the South, and by a population descendent from cosmopolitan *D. melanogaster* in the North [11,21,38]. Where these populations encountered each other in eastern North America, they form an ancestry cline where southern populations have a greater contribution of African ancestry than northern populations [21].

Previous work on these ancestry clines has shown that ancestry proportions vary across populations with increasing proportions of cosmopolitan alleles in more temperate localities. Evidence suggests spatially varying selection affects the distribution of genetic variants [40–45]. Furthermore, strong epistatic reproductive isolation barriers partially isolate individuals from northern and southern populations along this ancestry cline [46,47]. This may be generally consistent with recent observations of ancestry-associated epistatic fitness interactions within a *D. melanogaster* population in North Carolina [17], and with the observation of widespread fitness epistasis between populations of this species more generally [48]. There is therefore good reason to believe that natural selection has acted to shape LA clines that are tightly linked to selected mutations in these *D. melanogaster* populations.

Here, we show that African ancestry in North American *D. melanogaster* populations is negatively correlated with recombination rates, consistent with more efficient selection against foreign ancestry in high recombination rate regions of the genome. We also find that the X chromosome displays a higher rate of LA outlier loci, potentially consistent with a greater role of the X chromosome in clinal adaptation. Clinal loci are disproportionately likely to be associated with high level gene regulatory protein complexes, and may play important roles in ecological divergence between African and Cosmopolitan *D. melanogaster* populations. Furthermore, we identify numerous loci with decreased African ancestry across all populations, which suggests that these alleles that are disfavored on predominantly cosmopolitan genetic backgrounds. This subset of loci is enriched for genes related to oogenesis, potentially consistent with epistatic interactions that affect female reproductive success in these populations.

## Results and Discussion

### The Model

Although admixed populations often are diploid, we derived a general model of ploidy in which the individual has *n* gene copies at each locus, i.e. for diploid species *n* = 2. In practice, sequences are often obtained from fully or partially inbred individuals (e.g. [39,49]), which represent only a single uniquely derived chromosome. It is also common to pool individuals prior to sequencing for allele frequency estimation, so called pool-seq (e.g. [21,40,42,50–53]). If the pooling fractions are exactly equal, such a sample of *b* diploid individuals can be treated as a sample from a single individual with ploidy *n* = 2*b*. Although that requirement is restrictive, pool-seq has been experimentally validated as a method for accurate allele frequency estimation—*i.e.* alleles are approximately binomially sampled from the sample allele frequencies [54]. We therefore aimed to derive a model that can accommodate arbitrary sample ploidies. In the model, we assumed that the focal population was founded following a single discrete admixture event between two ancestral subpopulations, labeled 0 and 1, with admixture proportions 1-*m* and *m*, respectively, at a time *t* generations in the past. We modeled emission probabilities such that the method can work directly on read pileup data, rather than high quality known genotypes. Briefly, in our model, we specify an HMM {*H*_*v*_} with state space *S* = {0,1,…,*n*}, where *H*_*v*_ = *i*, *i* ∈ *S*, indicates that in the *v*th position *i* chromosomes are from population 0 and *n − i* chromosomes are from population 1. In other words, this HMM enables one to estimate what ancestry frequencies are present at a given site along a chromosome within a sample. Importantly, we designed this method to simultaneously estimate the time of admixture, which is related to the correlation between ancestry informative markers along a chromosome. See Methods for a complete description of the HMM including the emissions and transition probability calculations. The source code and manual are available at https://github.com/russcd/Ancestry_HMM. For this model, it is assumed that the number of chromosomes present in a sample, *n*, is known and that the global ancestry proportion, *m*, is known. As there are many methods for accurately estimating *m* in a wide variety of contexts implemented in standard population genetic analysis pipelines [19,20], we believe this assumption is not too restrictive.

### Admixture Simulation Framework

In order to test our method with data of known provenance, we also developed an approach for simulating chromosomes sampled from admixed populations. Briefly, we first simulated genetic diversity consistent with ancestral populations using a coalescent simulation method [55]. We then generated ancestry tracts consistent with admixture models developed to test our inference method using the forward-time admixture simulation program, SELAM [56]. We retained a portion of each coalescent-generated population to serve as a reference panel for allele frequency and LD estimation. We then took the remaining chromosomes and placed them on the appropriate ancestry tracts in admixed chromosomes. Finally, we generated read counts for these chromosomes, or pools of chromosomes for samples with ploidy greater than one, via binomial sampling from the genotype frequencies of the sample. Implicitly, this procedure assumes that the allele frequencies in the reference panel and the admixed individuals whose ancestry is from a given reference panels are equivalent. For large, well-mixed populations such as those of *D. melanogaster*, this is likely to be a reasonable assumption. Nonetheless, below we assess the impact of differences in the ancestral allele frequencies for plausible demographic models in this species.

### Dependence on Ancestral Linkage Disequilibrium

Within an admixed population, there are two sources of LD. LD that is induced due to the correlation of alleles from the same ancestry type (*i.e.* admixture LD), and LD that is present within each of the ancestral populations (ancestral LD). Admixture LD, is the signal of LA that we seek to detect using the HMM. The second type, ancestral LD, limits the independence of the ancestral information captured by each marker, and is expected to confound HMM-based analyses, particularly as we aimed to estimate the time since admixture within this framework. We therefore sought to quantify the effect of ancestral LD by discarding one of each pair of sites in LD within either ancestral population. We found that ancestral LD tends to increase admixture time estimates obtained using our method, and we decreased the cutoff of the LD parameter, |*r*|, by 0.1 until the time estimates obtained for single chromosomes were unbiased with respect to the true time since admixture. We found that |*r*| ≤ 0.4 fit this criterion, although for relatively ancient admixture events with highly skewed ancestry proportions—*i.e. m* < 0.1 or *m* > 0.9—some residual bias was apparent in the estimates of admixture time (Figure 1). This reflects the fact that the SMC’ ancestry tract distribution performs poorly with highly skewed ancestry proportions and especially for long times since admixture [57].

**Figure 1.**
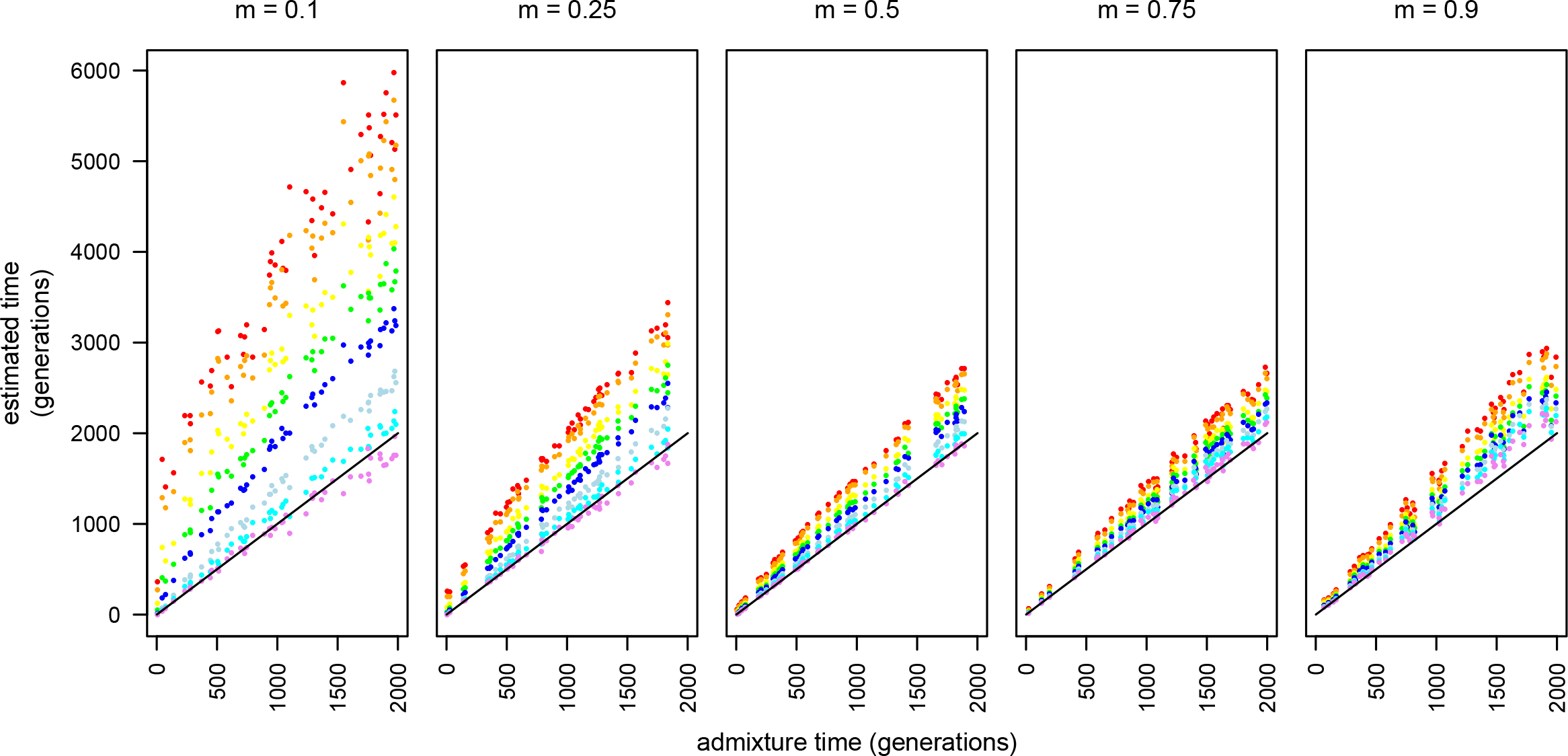
The effect of increasing stringency with ancestral LD pruning. From left to right, ancestry proportions are 0.1, 0.25, 0.5, 0.75 and 0.9. |r| cutoffs are: none (red), 1.0 (orange), 0.9 (yellow), 0.8 (green), 0.7 (dark blue), 0.6 (cyan), 0.5 (indigo), and 0.4 (violet). The solid line indicates the expectation for unbiased time estimation. All read data were simulated with ploidy = 1. True admixture time was drawn from a uniform (0, 2000) distribution.

Figure 1 also reveals a striking difference between otherwise equivalently skewed admixture proportions. For example when *m* = 0.1, there was a much larger effect of ancestral LD than when *m* = 0.9. This is due to differences in the variability of LD within the ancestral populations. That is, due to the strong population bottleneck, cosmopolitan *D. melanogaster* populations have substantially more LD and fewer polymorphic sites than African *D. melanogaster* populations. Because the time estimation procedure appears to be sensitive to the amount of ancestral LD present in the data, simulations of the type we described here may be necessary to determine what |*r*| cutoffs are required to produce unbiased time estimates given the ancestral LD of the populations in a given analysis using this method.

### Accuracy and Applications to Diploid and Pooled Samples

We next sought to quantify the accuracy of our approach across varying sample ploidies and times since admixture (Figure 2). Especially for moderate and short admixture times (*i.e.* 0—500 generations), our method performed well for all ploidies considered and we were able to accurately recover the correct admixture time with relatively little bias. However, as true admixture time increases, the time estimates for pooled samples become significantly less reliable and show a clear negative bias. Nonetheless, across the range of times presented in Figure 2, samples of ploidy one and two showed little bias, and we therefore believe our method will produce sufficiently accurate admixture time estimates for a wide variety of applications.

**Figure 2.**
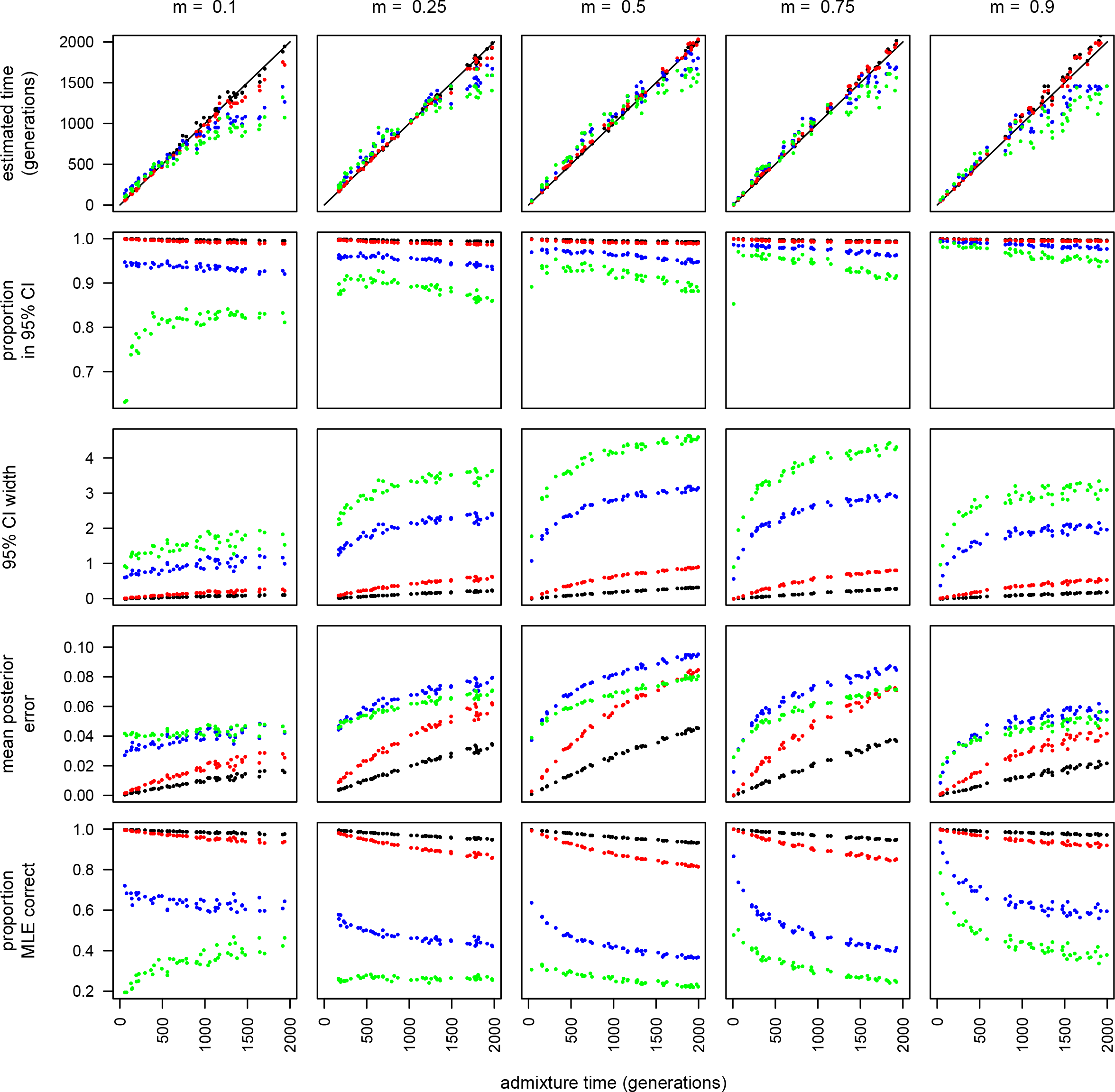
Time estimates and accuracy statistics for samples of varying ploidies. From left to right, ancestry proportions are 0.1, 0.25, 0.5, 0.75 and 0.9. Each sample ploidy is represented by one point color with ploidy one (black), two (red), ten (blue) and twenty (green). From top to bottom, each row is the estimated time in generations, the proportion of sites where the true state is within the 95% credible interval, the width of the 95% credible interval, the mean posterior error, and the proportion of sites where the maximum likelihood estimate is equal to the true state.

All measures of accuracy decrease with increasing time since admixture (Figure 2). However, even for relatively long times since admixture—2000 generations—and for large sample ploidies, the mean posterior error remained relatively low for all ancestry proportions and for long times since admixture. This indicates that this approach may be sufficiently accurate for a wide variety of applications, sequencing depths, and sample ploidies. Nonetheless, the proportion of sites within the 95% credible interval decreased with larger pool sizes and it is clear that for larger pools the posterior credible interval tends to be too narrow. Therefore, correcting for this bias may be necessary for applications that are sensitive to the accuracy of the credible interval.

An important consideration is that estimates of *t* will be reliable only if the local recombination rates are known with reasonably high accuracy [58]. In many species, an accurate broad-scale map is available. However, fine-scale variation in recombination rates has only been documented for a few model species. Therefore, for relatively short to moderate times since admixture, error in the genetic map is expected to have a limited impact on date estimates. However, for longer times since admixture, this factor has the potential to bias estimates of *t* [58], particularly in species with large variance in local recombination rates (*e.g*. due to hotspots). Since *D. melanogaster* has one of the best recombination maps currently available in any species [59] and because we do not aim to estimate time in our applications, we do not believe this will heavily impact the analyses we present below. However, for most applications, it will be necessary to consider the impact of error in the assumed genetic map to accurately interpret estimates of t obtained using this method. We emphasize that this challenge is not unique to this application, but will impact virtually all ancestry estimation methods that rely on a genetic map for estimating the time since admixture.

### Non-Independence Among Ancestry Tracts

As described above, estimates of the time of admixture demonstrate an apparent bias in pools of higher ploidy (Figure 2). Specifically, time tends to be slightly overestimated for relatively short admixture times and underestimated at relatively long admixture times. This is particularly apparent at highly skewed ancestry proportions. Given that this bias is primarily evident in pools of 10 to 20 individuals, we hypothesized that it might be due to the non-independence of ancestry tracts among chromosomes, which should tend to disproportionately affect samples of higher ploidy because all ancestry breakpoints are assumed to be independent in our model. To test this, we simulated genotype data from independent and identically distributed exponential tract lengths as is assumed by our model. When we ran our HMM on this dataset, we found that no bias is evident for simulations of up to 2000 generations (Supplemental Figure S1), indicating that the primary cause of this bias was violations in the real data of the independence of ancestry tracts that we assumed when computing the transition probabilities. However, it should be possible to quantify and correct for this bias in applications of this method that aim to estimate the time since admixture.

### Robustness to Unknown Population Size

The transition probabilities of this HMM depend on knowledge of the population size. In practice, this parameter is unlikely to be known with certainty. Hence, to assess the impact of misspecification of the population size, we performed simulations using a range of population sizes that span three orders of magnitude (*N* = 100, 1000, 10000, and 100000). All analyses presented here were conducted by applying our HMM to haploid and diploid samples, but qualitatively similar results hold for samples of larger ploidy. We then analyzed these data assuming the default population size, 10000, is correct. For relatively short times since admixture, there was not a clear bias for any of the true population sizes considered. However, at longer true admixture times, estimated admixture times for both *N=*100 and *N=*1000 asymptote at a number of generations near to the population sizes. This result reflects the fact that smaller populations will tend to coalesce at a portion of the loci in the genome relatively quickly, and ancestry tracts cannot become smaller following coalescence. Nonetheless, the accuracy of LAI remained high even when time estimates were unreliable (Supplemental Figure S2) for the tested marker densities and patterns of LD. Furthermore, in some cases it should be straightforward to determine if a population has coalesced to either ancestry state at a large portion of the loci in the genome, potentially obviating this issue.

A more subtle departure from the expectation was evident for population sizes that are larger than we assumed in analyzing these data (Supplemental Figure S2). This likely reflects the fact that the probability of back coalescence to the previous marginal genealogy to the left after a recombination event is inversely related to the population size. Hence, the rate of transition between ancestry types is actually slightly higher in larger populations where back coalescence is less likely than we assumed during the LAI procedure. This produced a slight upward bias in the estimates of admixture time when the population was assumed to be smaller than it is in reality. However, this bias appears to be relatively minor, and we expect that time estimates obtained using this method will be useful so long as population sizes can be approximated to within an order of magnitude. Of course, this bias is not unique to our application, and it will affect methods that aim to estimate admixture time after LAI as well. That is, estimating the correct effective population size is an inherent problem for all admixture demographic inference methods.

### Application to Ancient Admixture

Although it is clear that accurately estimating relatively ancient admixture times is challenging in higher ploidy samples, we sought to determine the limits of our approach for LAI and time estimation for longer admixture times for haploid sequence data. Because of rapid coalescence in smaller samples (see above), we performed admixture simulations with a diploid effective population size of 100,000. It is clear that there is a limit to the inferences that can be made directly using our method. Like the higher ploidy samples, time estimates for haploid samples departed from expectations shortly after 2,000 generations since admixture (Supplemental Figure S3). Nonetheless, the magnitude of this bias is slight, and it is likely that it could be corrected for when applying this method even for very ancient admixture events. For all admixture times considered, LAI remained acceptably accurate despite the slight bias in time estimates (Supplemental Figure S3).

### Reference Panel Size

One question is what effect varying the reference panel sizes will have on LAI inference using this method. We therefore compared results from reference panels of size 10 chromosomes with those from panels of size 100 chromosomes (Supplemental Figure S4). As with results obtained for reference panels of size 50, panels of size 100 were sufficient to accurately estimate admixture time and LA over many generations since admixture. Whereas, when panel sizes were just 10 chromosomes, time estimates were clearly biased and the result was variable across ancestry proportions (Supplemental Figure S4). However, since there was a strong correlation between true and estimated admixture times even with relatively small panel sizes, it may therefore be possible to infer the correct time by quantifying this bias through simulation and correcting for it. Furthermore, although LAI is clearly less reliable with smaller panels, these results are not altogether discouraging and this approach, in conjunction with modest reference panels may still be effective for some applications.

### Allele Frequency Differences Between Ancestral and Admixed Populations

Ultimately, there are three reasons why allele frequencies in the reference panels and in the admixed population panel would be expected to differ beyond that expected from binomial samples with the same mean. First, some amount of genetic drift may have occurred in the ancestral population and in the admixed population in the time since the admixed population was founded. Second, in some cases, it is infeasible to sample the ancestral population of an admixed group, and a genetically divergent population must suffice as the reference panel if this method is to be used. Third, divergent selection may quickly modify allele frequencies between admixed and ancestral populations. Hence, genetic divergence between reference and admixed populations may be an important challenge for this method.

To address this, we simulated the second scenario, where increasingly divergent populations are used as the reference panels to study admixed populations. In order to make this relevant to the application to *D. melanogaster* populations, below, we selected times for divergence that might be consistent with differences across continental populations in Sub-Saharan Africa and in Cosmopolitan populations. Although time estimates obtained using this approach are weakly positively biased with increasing divergence between the ancestral population and reference panels, the accuracy of this LAI method is largely unaffected (Supplemental Figure S5). Hence, for biological scenarios potentially consistent with those of *D. melanogaster* ancestral populations, we do not expect this challenge to strongly bias our method. Nonetheless, in applications to other populations, with potentially differently structured ancestral populations, it would be necessary to examine the effects of this bias in detail.

### High Sample Ploidy

In a wide variety of pool-seq applications, samples are pooled in larger groups than we have considered above (*e.g.* [40,50,52]). We are therefore interested in determining how our method will perform on pools of 100 individuals. Towards this, we performed simulations as before, but we designed our parameters to resemble those of the pooled sequencing data that we analyze in the application of this method below. Specifically, we simulated data with a mean sequencing depth of 25, a time since admixture of 1500 generations, and an ancestry proportion of 0.8. Consistent with results for ploidy 20, we found that time tends to be dramatically underestimated (*i.e.* the mean estimate of admixture time was 680 generations). However, when we provided the time since admixture, our method produced reasonably accurate LAIs for these samples. Although the posterior credible interval was again too narrow, the mean posterior error was just 0.053 when expressed as an ancestry frequency, indicating that this approach can produce LA estimates that are close to their true values for existing sequencing datasets (*e.g*. Figure 3). However, the HMM’s run time increases dramatically for higher ploidy samples and higher sequencing depths, a factor that may affect the utility of this program for some analyses. Nonetheless, for more than 36,000 markers, a sample ploidy of 100 and a mean sequencing depth of 25, the average runtime was approximately 42 hours. In contrast, for the same set of parameters, but where individuals are sequenced and analyzed as diploids, the mean runtime was just 8 minutes (See Supplemental Table S1 for a comparison of run times across many parameter sets).

**Figure 3.**
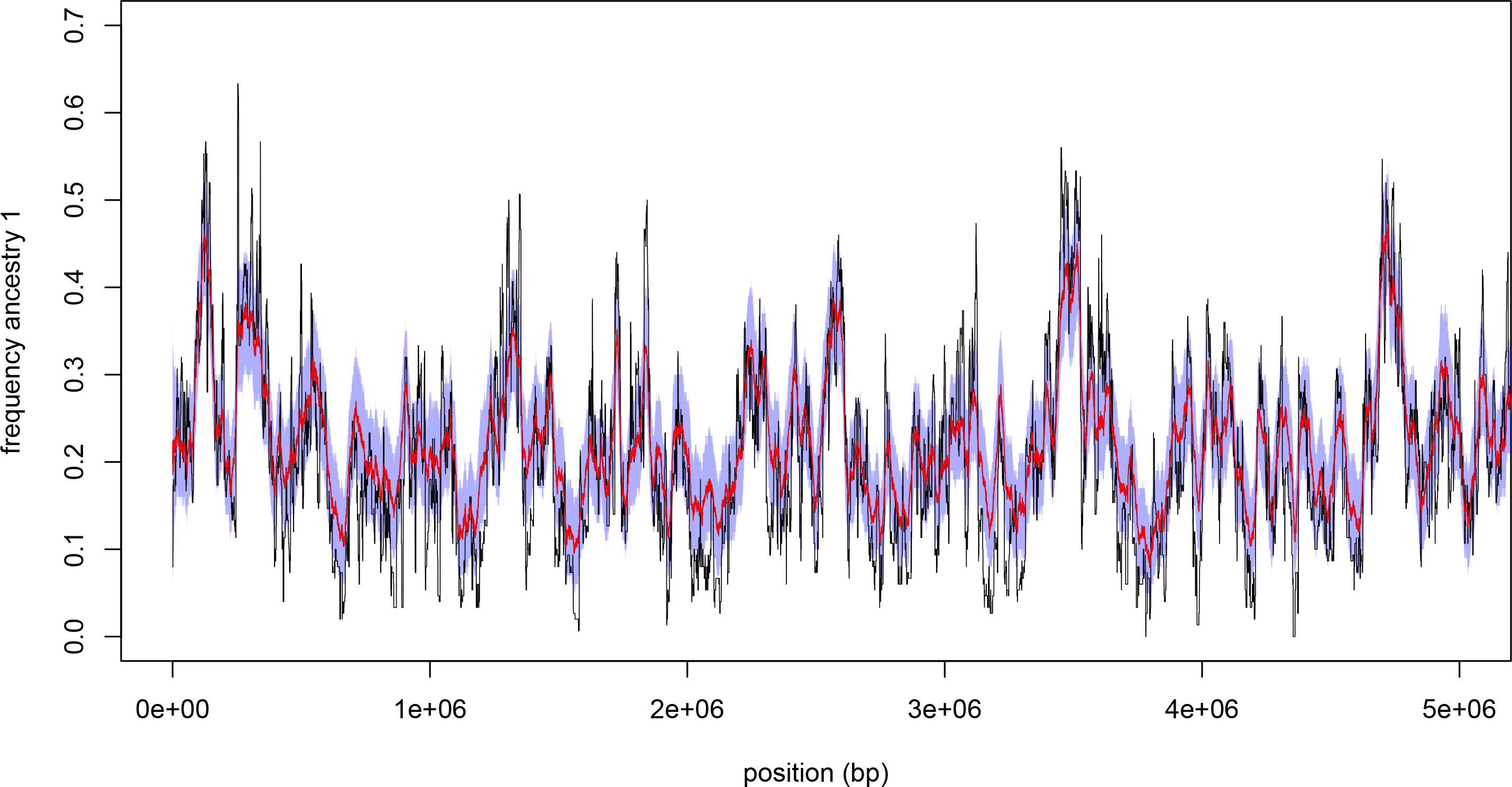
Accuracy of the HMM for samples of high ploidy. The 95% credible interval (shaded blue region), and the posterior mean (red) contrasted with the true ancestry frequencies (black). Simulated data were generated with an admixture time of 1500 generations, an ancestry proportion of 0.2, a sample ploidy of 100, and a mean sequencing depth of 25.

### Robustness to Deviations From the Neutral Demographic Model

An important concern is that many biologically plausible admixture models would violate the assumptions of this inference method. In particular, continuous migration and selection acting on alleles from one parental population are two potential causes of deviation from the expected model in the true data. To assess the extent of this potential bias, we performed additional simulations. First, we considered continuous migration at a constant rate that began *t* generations prior to sampling. In simulations with continuous migration, additional non-recombinant migrants enter the population each generation. Relative to a single pulse admixture model, this indicates that the ancestry tract lengths will tend to be longer than those under a single pulse admixture model in which all individuals entered at time *t*. Indeed, we found that admixture times tended to be underestimated with models of continuous migration. However, the accuracy of LAI remained high across all situations considered here (Table 1), indicating that the LAI aspect of this approach may be robust to alternative demographic models.

**Table 1.**
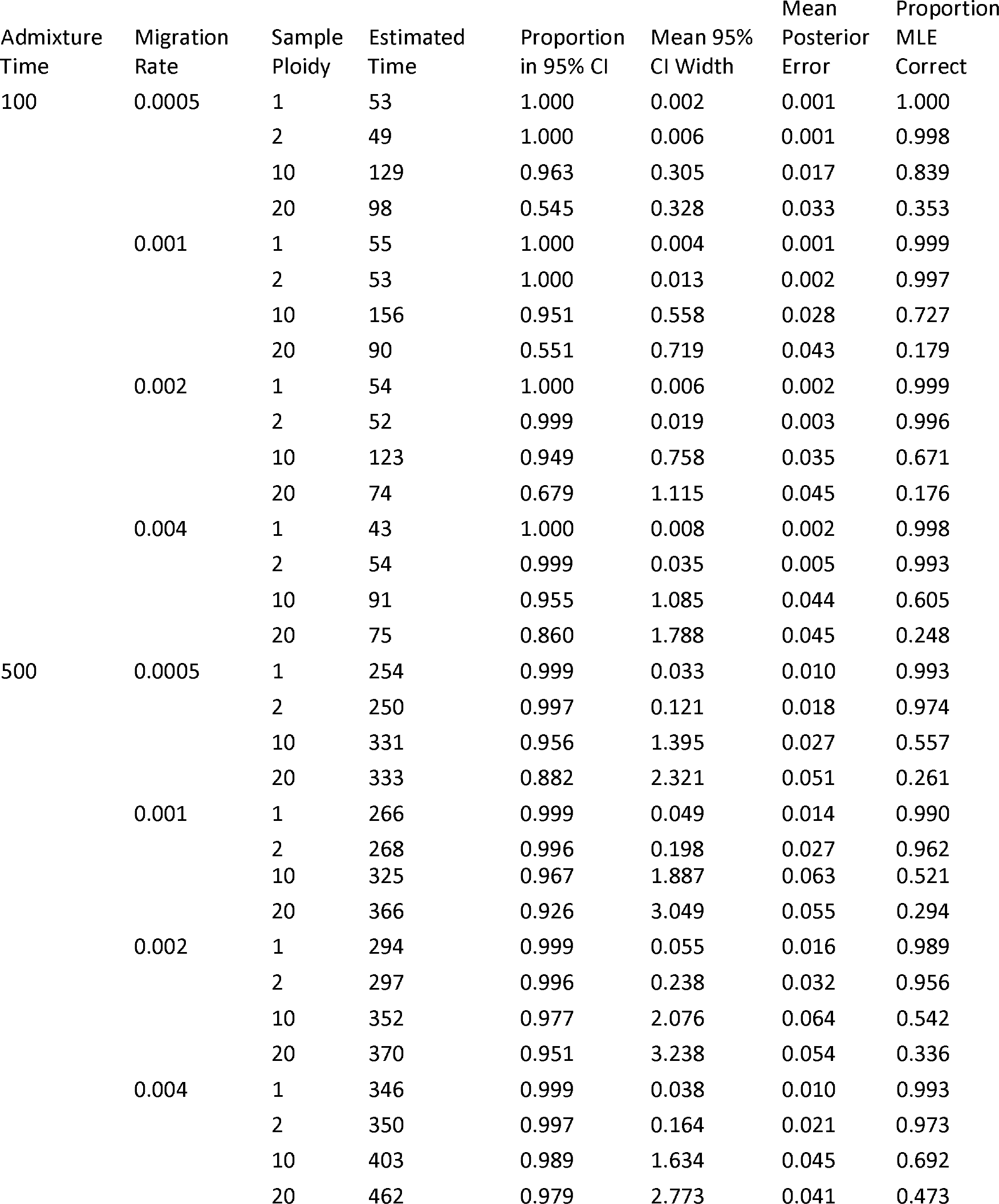
Parameter estimation and LAI when admixture occurs at a constant rate, rather than in a single pulse.

In the second set of simulations, we considered additive selection on alleles that are perfectly correlated with local ancestry in a given region (*i.e.* selected sites with frequencies 0 in population 0 and frequency 1 in population 1), and experience relatively strong selection (selective coefficients were between 0.005 and 0.05). We placed selected sites at 2, 5, 10 and 20 loci distributed randomly across the simulated chromosome, where admixture occurred through a single pulse. Ancestry tracts tend to be longer immediately surrounding selected sites, and we therefore expected admixture time to be underestimated when selection is widespread. When the number of selected loci was small, time estimates were nearly unbiased (Table 2), suggesting that our approach can yield reliable admixture time estimates despite the presence of a small number of selected loci (*i.e.* 2 selected loci on a chromosome arm). However, with more widespread selection on alleles associated with local ancestry, time estimates showed a downward bias that increased with increasing numbers of selected loci. This is likely because selected loci will tend to be associated with longer ancestry tracts due to hitchhiking. However, the accuracy of the LAI remains high for all selection scenarios that we considered here, further indicating that our method can robustly delineate LA, even when the data violate assumptions of the inference method (Table 1,2).

**Table 2.**
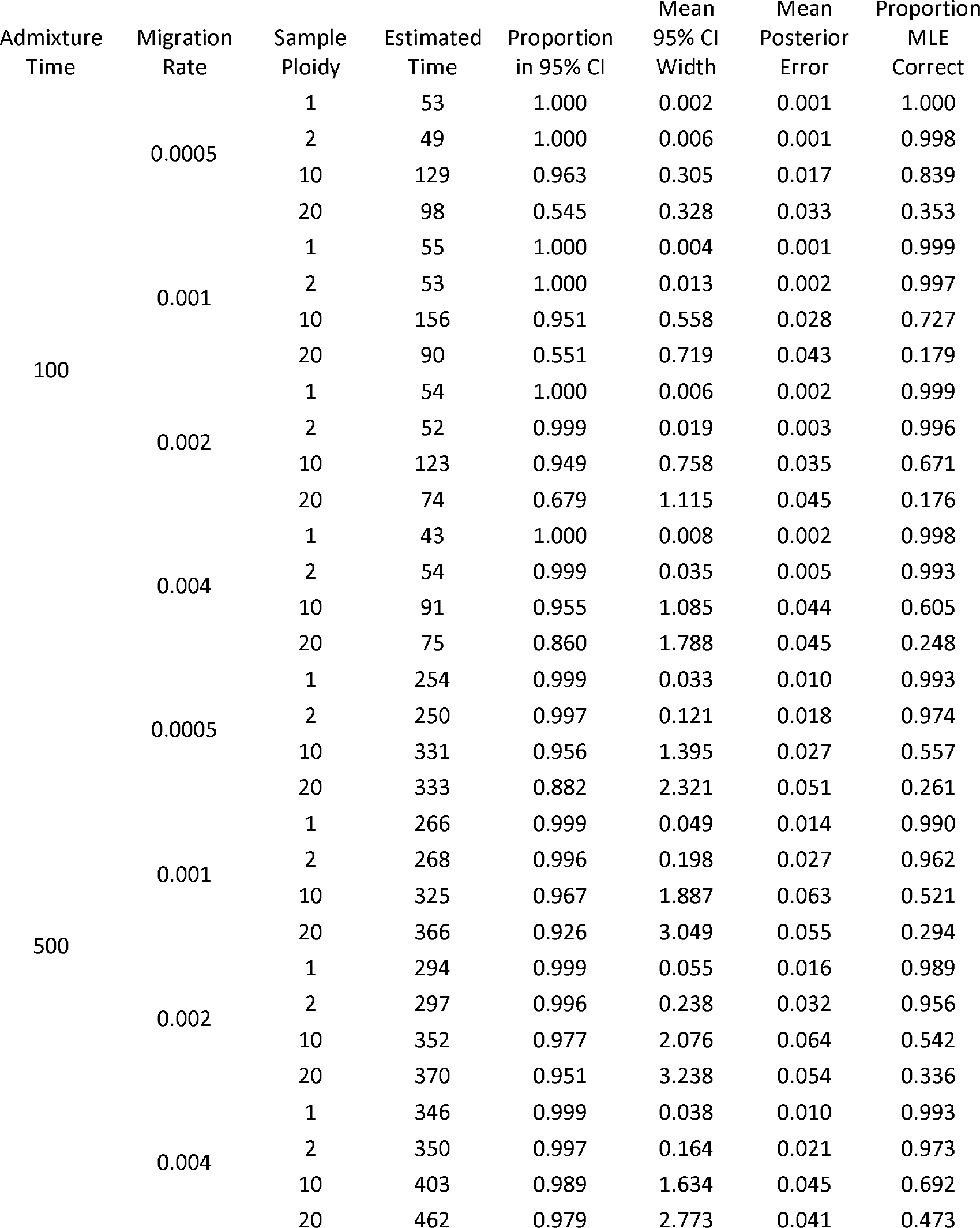
Parameter estimation and LAI when a subset of loci experience natural selection in the admixed population.

### Comparison to WinPop

We next compared the results of our method to those of WinPop [34]. Because WinPop accepts only diploid genotypes, we provided this program diploid genotype data. However, for these comparisons, we still ran our method on simulated read pileups with the mean depth equal to 2. WinPop was originally designed for local ancestry inference in very recently admixed populations. As expected, WinPop performed acceptably for very short admixture times, but rapidly decreased in performance with increasing time (Supplemental Figure S6). However, by default, WinPop removes sites in strong LD within the admixed samples, which includes ancestral LD, but also admixture LD—the exact signal LAI methods use to identify ancestry tracts.

We therefore reran WinPop, but instead of pruning LD within the admixed population, we removed sites in strong LD within the ancestral populations as described above in our method. With this modification, WinPop performs nearly as well as our method, but remains slightly less accurate especially at longer admixture times (Supplemental Figure S6). This difference presumably reflects the windowed-based approach of WinPop. At longer times since admixture a given genomic window may overlap a breakpoint between ancestry tracts. Although the performance is nearly comparable with this modification, we emphasize that our method enables users to estimate the time since admixture, where this must be supplied for WinPop, and allows for LAI on read pileups, therefore incorporating genotype uncertainty into the LAI procedure. Indeed our method is more accurate at longer timescales even when supplied with considerably lower quality read data. However WinPop supports LAI with multiple ancestral populations, which our method currently does not (but see Conclusions). Furthermore many LAI algorithms utilize haplotype information, which may be particularly valuable in populations where LD extends across large distances as in *e.g*. human populations.

### Assessing Applications to Human Populations

Given the strong interest in studying admixture and local ancestry in human populations (*e.g*. [22–25]), it is useful to ask if our method can be applied to data consistent with admixed populations of humans. Towards that goal, we simulated data similar to what would be observed in admixture between modern European and African lineages and applied our HMM to estimate admixture times and LA. We found that our method can accurately estimate admixture times for relatively short times since admixture, however, substantially more stringent LD pruning in the reference panels is necessary to produce unbiased estimates (Figure 4). This may be expected given that linkage disequilibrium extends across longer distances in human populations than it does in *D. melanogaster*. In other words, the scales of ancestral LD and admixture LD become similar rapidly in admixed human populations. Furthermore, this approach yields accurate time estimates for shorter times since admixture than with genetic data consistent with *D. melanogaster* populations. For a relatively short time since admixture, around 100 generations, it is possible to obtain accurate and approximately unbiased estimates of the admixture time over a wide range of ancestry proportions, indicating that this method may be applicable to recently admixed human populations as well (Figure 4). Nonetheless, this result underscores the need to examine biases associated with LD pruning in this approach prior to application to a given dataset.

**Figure 4.**
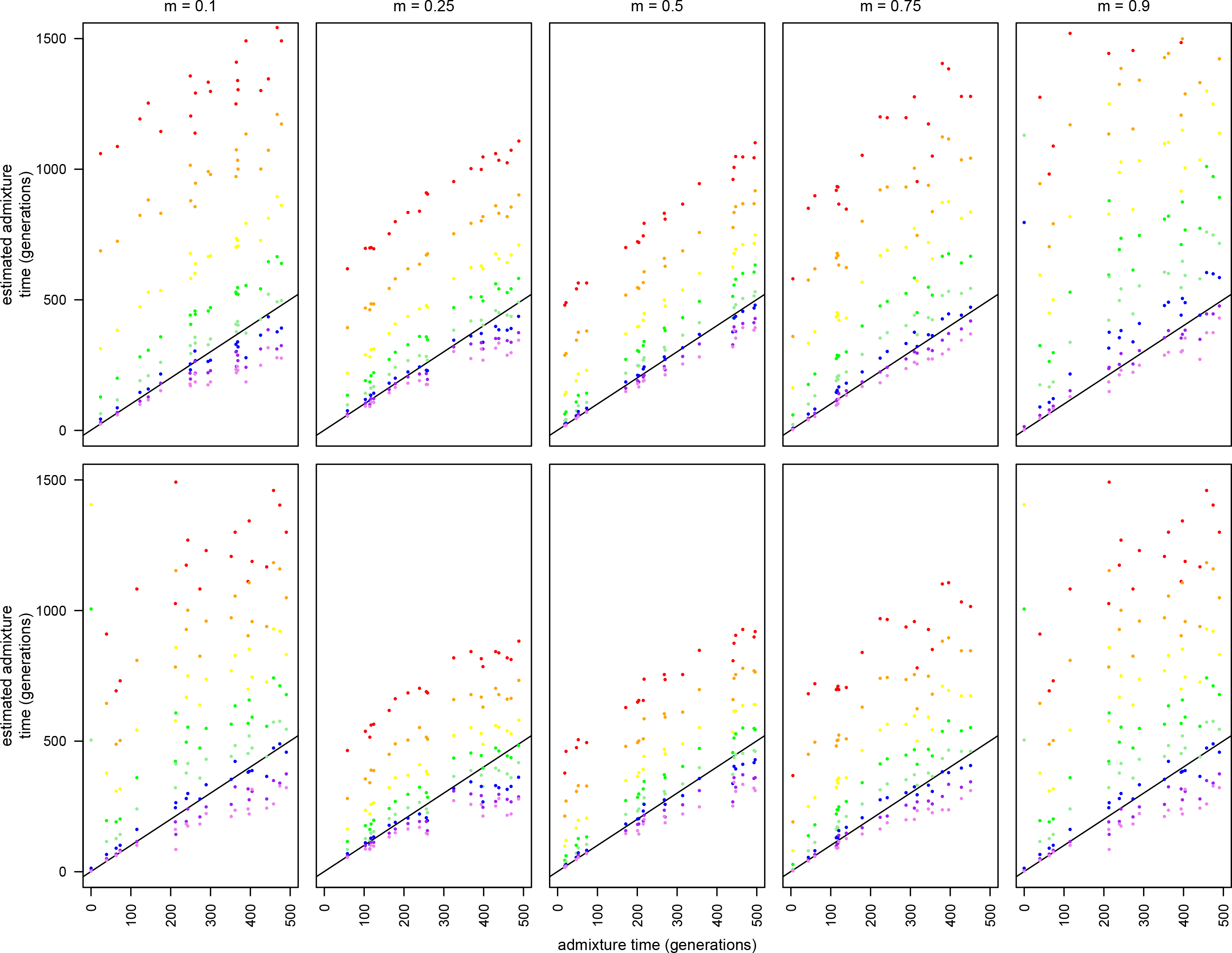
Admixture time estimates for simulated data consistent with variation present in modern European and African populations. From left to right, *m* = 0.1, *m* = 0.25, *m* = 0.5, *m* = 0.75, *m* = 0.9. The top row is completely phased chromosomes and the bottom row is unphased diploid data.

### Bias in LAI due to Uncertainty in Time of Admixture

To demonstrate that assumptions about the number of generations since admixture have the potential to bias LAI, we analyzed a SNP-array dataset from Greenlandic Inuits [60,61]. The authors had previously noted a significant impact of *t* on the LAI results produced using RFMix [24], which we were able to reproduce here for chromosome 10 (Supplemental Figure S7). Indeed, even for comparisons between *t* = 5 and *t* = 20, both of which may be biologically plausible for these populations, the mean difference in posterior probabilities between samples estimated using RFMix was 0.0903 (Supplemental Figure S7). However, when we applied our method to these data, a clear optimum from *t* was obtained at approximately 6–7 generations prior to the present, which is close to the plausible times of admixture for these populations (Supplemental Figure S7). This comparison therefore demonstrates that even relatively minor changes in assumptions of *t* have the potential to strongly impact LAI results, and underscores the importance of simultaneously performing LAI while estimating *t*.

However, these results also indicate that our method may not be robust in situations where the background LD is high and ancestry informative markers are neither common nor distributed evenly across the genome. When we compared the results of our method at *t* = 5 and at *t* = 20, we also obtained differences in the mean posterior among individuals as with RFMix. However, one notable difference is that the mean posterior difference using RFMix has a particularly high variance and therefore higher mean error (Supplemental Figure S7), but actually a lower median difference than we found using our method. There are likely two causes for differences observed in the mean ancestry posterior among individuals. First, the datasets considered were generated with a metabochip SNP-chip [62], which contains a highly non-uniform distribution of markers across the genome. Second, the ancestral LD in the Inuit population is extensive [61], and we could only retain a relatively small proportion of the markers after LD pruning in the reference panels. These results therefore also underscore the challenges of LAI when the signal to noise ratio is low as may be the case in some human populations, for which LD is extensive, and for some sequencing strategies.

### Bias due to Incorrect Estimates of *t* and *m*

Although in general it is straightforward to estimate *m* from genome-wide data, in some cases this parameter may be misestimated prior to LAI. We therefore sought to quantify this potential effect by performing LAI after supplying incorrect values of m. In general, we found that values close to the true range, *i.e.* within 0.05 of the true *m*, tend to yield reasonably accurate time estimates. However, increasingly incorrect values produce sharply downwardly biased time estimates and this effect is especially pronounced for highly skewed true *m* (Supplemental Figure S8). As could be expected given the robustness of LAI to many perturbations (above), when the incorrect *t* is supplied to the program, the LA results remain reasonable. However it is worth noting that the penalty appears to be greatest when *t* is too small rather than too large (Supplemental Figure S9).

### Estimating Confidence Intervals for *t*

Although this is not a primary focus for this work, for some users it may be of interest to construct confidence intervals for estimates of *t*. We recommend the block bootstrap as the preferred method for estimating confidence interval for *t*, and we have written a script that will produce these (available on the github page for this project: https://github.com/russcd/Ancestry_HMM). Simulations confirm that this can produce confidence intervals overlapping the true *t* (Supplemental Figure S10), but bias in *t* estimates for higher ploidy samples may still be apparent in some cases.

### Patterns of LA on Inversion Bearing Chromosomes in *D. melanogaster*

Given their effects suppressing recombination in large genomic regions, chromosomal inversions may be expected to strongly affect LAI [2,63]. Although we attempted to limit the impact of chromosomal inversions by eliminating known polymorphic arrangements from the reference panels (see methods), many known inversions are present within the pool-seq samples we aimed to analyze [64]. We therefore focused on known inverted haplotypes within the DGPR samples [63,65–67], which are comprised of inbred individuals, and therefore phase is known across the entire chromosome.

In comparing LA estimates between inverted and standard arrangements, it is clear that chromosomal inversions can substantially affect LA across the genomes (Figure 5). In general, the chromosomal inversions considered in this work originated in African populations of *D. melanogaster* [63], and consistent with this observation, most inversion bearing chromosomes showed evidence for elevated African ancestry. This was particularly evident in the regions surrounding breakpoints, where recombination with standard arrangement chromosomes is most strongly suppressed. Importantly, this pattern continued outside of inversion breakpoints as well, consistent with numerous observations that recombination is repressed in heterokaryotypes in regions well outside of the inversion breakpoints in *Drosophila* (*e.g*. [2,63,68]). In(3R)Mo is an exception to this general pattern of elevated African ancestry within inverted arrangements (Figure 5). This inversion originated within a cosmopolitan population [63], and has only rarely been observed within sub-Saharan Africa [69,70]. Consistent with these observations, In(3R)Mo displayed lower overall African ancestry than chromosome arm 3R than standard arrangement chromosomes.

**Figure 5.**
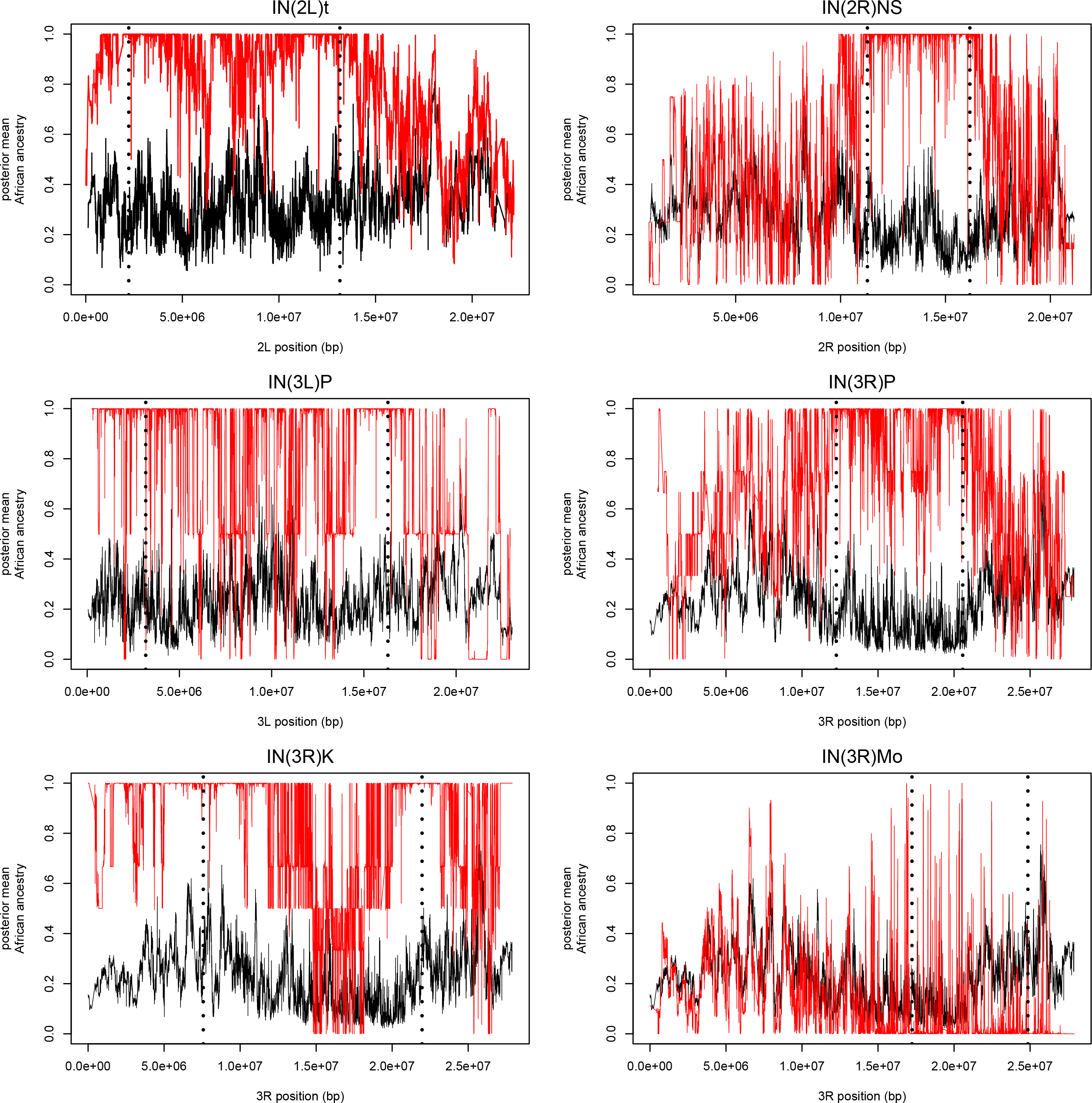
Local ancestry of inversion bearing chromosomes (red) compared with those of standard arrangement chromosomes (black) for the same chromosome arm. Positions of inversion breakpoints, as reported in [63,67] are shown as vertical dashed lines.

Although chromosomal inversions may affect patterns of LA in the genome on this ancestry cline, we believed including chromosomal inversions in the pool-seq datasets would not heavily bias our analysis of LA clines. Inversions tend to be low frequency in most populations studied [64], and because they affect LA in broad swaths of the genome—sometimes entire chromosome arms—including inversions is unlikely to affect LA cline outlier identification which appears to affect much finer scale LA (below). Furthermore, inversion breakpoint regions were not enriched for LA cline outliers in our analysis (Supplemental Table S2), suggesting that inversions have a limited impact on overall patterns of local ancestry on this cline. Nonetheless, the LAI complications associated with chromosomal inversions should be considered when testing selective hypotheses for chromosomal inversions as genetic differentiation may be related to LA, rather than arrangement-specific selection in admixed populations such as those found in North America.

### Application to *D. melanogaster* Ancestry Clines

Finally, we applied our method to ancestry clines between cosmopolitan and African ancestry *D. melanogaster*. Genomic variation across two ancestry clines have been studied previously [21,38,40,52]. In particular, the cline on the east coast of North America has been sampled densely by sequencing large pools of individuals to estimate allele frequencies, and previous work has shown that the overall proportion of African ancestry is strongly correlated with latitude [21]. Consistent with this observation, we found a significant negative correlation for all chromosome arms between proportion of average African ancestry and latitude (rho = −0.891, −0.561, −0.912, −0.913, and −0.755, for 2L, 2R, 3L, 3R, and X respectively).

Although global ancestry proportions have previously been investigated in populations on this ancestry cline [21,38], these analyses neglected the potentially much richer information in patterns of LA across the genome. We therefore applied our method to these samples. Because of the relatively recent dual colonization history of these populations and subsequent mixing of genomes, a genome-wide ancestry cline is expected [21]. However, loci that depart significantly in clinality from the genome-wide background levels may indicate that natural selection is operating on a site linked to that locus.

### LA is Correlated with Recombination Rate

Previously Pool (2015) found that regions of low recombination are disproportionately enriched for African ancestry in the Raleigh, NC population [17]. Here, we find a similar pattern and we further find that is replicated across all populations that were assayed on this ancestry cline. Specifically, in all populations studied the proportion of African ancestry is significantly negatively correlated with local recombination rates (Figure 6). Ultimately, this correlation may have two causes. First, if selection is more efficient at purging African alleles in high recombination regions, these loci will tend to be removed preferentially in those genomic regions. An alternative explanation is that introgressing African alleles that are favored by selection would tend to bring larger linkage blocks along with them in the predominantly low recombination regions. Regardless of the specific source of natural selection, a neutral admixture model would not predict this robust correlation between LA and recombination rates within all populations, indicating that natural selection has played an important role in shaping LA on this ancestry cline.

**Figure 6.**
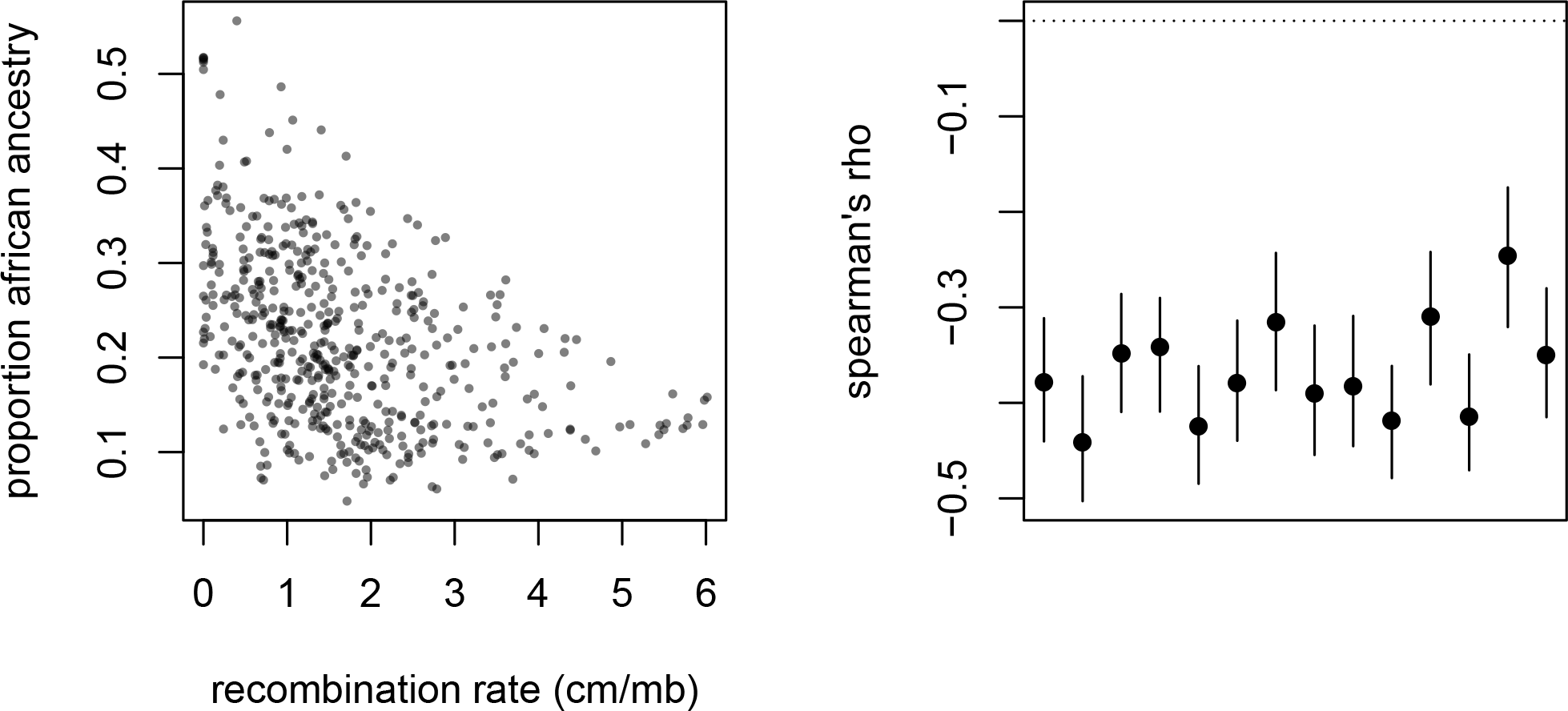
The relationship between the proportion of African ancestry proportion and local recombination rates in 100 ancestry informative SNP windows within the Raleigh, NC population (left). The correlation between the proportion of African ancestry proportion and local recombination rates in 100 ancestry informative SNP windows in all populations assayed (right). Lines indicate the 95% confidence interval obtained via block bootstrap replicates (see Methods).

### Robustness of LAI to Genomic Heterogeneity

Previous studies have found that heterogeneity in the genome with respect to ancestry informative markers may impact the accuracy of LAI [71]. To assess this possibility, we computed the mean difference between posterior mean estimates for the two samples from Florida and between the two samples from Maine. Importantly, because these pooled samples were created using different isofemale lines [40], this is a conservative test of our method since there will be true biological differences as well as stochastic sequencing differences between replicates from each population. We found no correlation between the mean difference of the posterior means and local recombination rates (P = 0.2353 and P = 0.7529, Spearman’s rank correlation for Florida and Maine respectively), indicating that the correlation observed between local recombination rates and LA is unlikely to be an artifact of differential accuracy of LAI in different genomic regions. However, it should be acknowledged that in some genomic regions it maybe challenging to unambiguously infer LA [17,71].

### Outlier LA Clines

Selection within admixed populations may take several distinct forms. On the one hand, loci that are favorable in the admixed population—either because they are favored on an admixed genetic background, enhance reproductive success in an admixed population, or are favorable in the local environment—will tend to achieve higher frequencies, and we would expect these sites to have a more positive correlation with latitude than the genome-wide average. Conversely, loci that are disfavored within the admixed population may be expected to skew towards a more negative correlation with latitude.

Although it is not possible to distinguish between these hypotheses directly, a majority of evidence suggests that selection has primarily acted to remove African ancestry from the largely Cosmopolitan genetic backgrounds found in the Northern portion of this ancestry cline. First, abundant evidence suggests pre-mating isolation barriers between some African and cosmopolitan populations [72–74]. Second, there is strong post-mating isolation between populations on the ends of this cline [46,47]. Third, we report here a strong negative correlation between LA frequency and local recombination rates (above). Finally, circumstantially, the local environment on the east coast of North America is perhaps most similar to the environment of Cosmopolitan compared to African ancestral populations, which further suggests that Cosmopolitan alleles are likely favored through locally adaptive mechanisms. For these reasons, we therefore examined loci that are outliers for a negative partial correlation with latitude, as this is the expected pattern for African alleles that are disfavored in more temperate populations. In other words, the outlier regions show a significantly stronger negative correlation between local African ancestry and latitude than the chromosome arm does on average.

There is an ongoing debate about the relative merits of an outlier approach versus more sophisticated models for detecting and quantifying selection in genome-wide scans. We believe that the difficulties of accurately estimating demographic parameters for this ancestry cline make the outlier approach most feasible for our purposes. Using our outlier approach, we identified 80 loci that showed the expected negative partial correlation with latitude (Figure 7). Although the specific statistical threshold that we employed is admittedly arbitrary, given the strength of evidence indicating widespread selection on local ancestry in this species (above), we expected that the tail of the LA cline distribution would be enriched for the genetic targets of selection.

**Figure 7.**
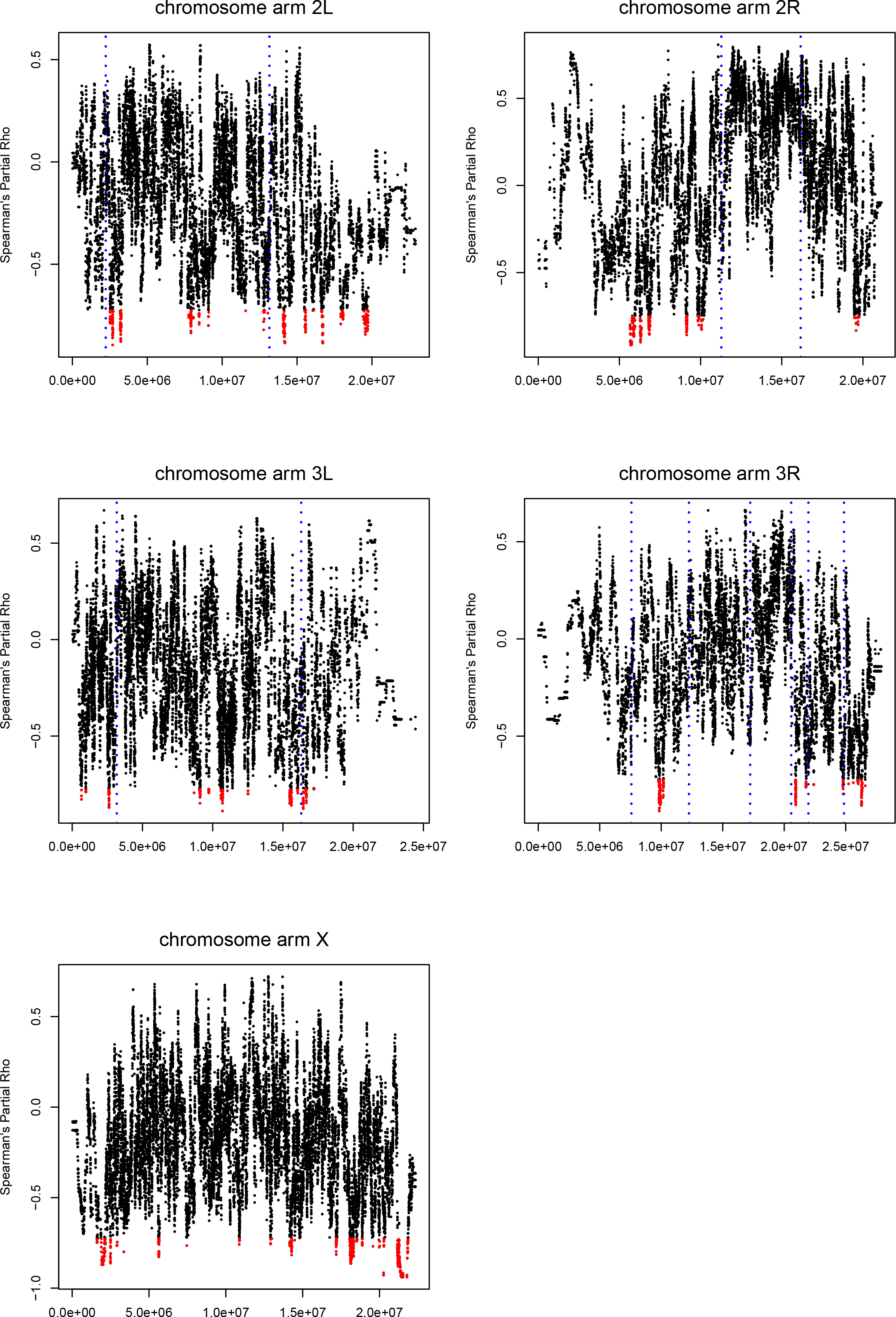
The partial correlation between LA and latitude with correction for chromosome-wide ancestry proportions. Sites for which the probability of the observed clinal relationship was less than 0.005 were retained as significant (red). Inversion breakpoints for inversions that are at polymorphic frequencies on this ancestry cline are shown as dotted blue lines.

### Differences Among Chromosome Arms

Due to the differences in inheritance, evolutionary theory predicts that selection will operate differently on the X chromosome relative to autosomal loci. Of specific relevance to this work, the large-X effect [75,76] is the observation that loci on the X chromosome contribute to reproductive isolation at a disproportionately high rate. Additionally, and potentially the cause of the large-X effect, due to the hemizygosity of X-linked loci, the X chromosome is expected to play a larger role in adaptive evolution, the so-called faster-X effect [77]. There is therefore reason to believe that the X chromosome will play a significant role in genetically isolating Cosmopolitan and African *D. melanogaster*.

Consistent with a larger role for the sex chromosomes in generating reproductive isolation or selective differentiation between *D. melanogaster* ancestral populations, we found that that the X chromosome has a lower mean African ancestry proportion than the autosomes in all populations. Furthermore, the X displays a stronger correlation between local recombination rates and the frequency of African ancestry than the autosomes in all 14 populations samples, potentially indicating that selection has had a disproportionately strong effect shaping patterns of local ancestry on this chromosome than on the autosomes. In addition, the X has a significantly higher rate of outlier LA clinal loci than the autosomes (23 LA outliers on the X, 57 on the Autosomes, p = 0.0341, one-tailed exact Poisson test). Although consistent with evolutionary theory, differences between autosomal arms and the X chromosome may also be explained in part by differences in effective recombination rates on the X chromosome than the autosomes, differences in power to identify LA clines associated with chromosome arm specific patterns, or by the disproportionately larger number of chromosomal inversions on the autosomes than on the X chromosome in these populations [64,69]. Distinguishing between this hypothesis and confounding factors will be central to determining whether key results from speciation research are replicated in much more recently diverged populations.

### Biological Properties of Outlier LA Clinal Loci

We next applied gene ontology analysis to the set of outlier genes to identify common biological attributes that may suggest more specific organismal phenotypes underlying LA clinal outliers. In total, we identified seven GO terms that remained significant after applying a 5% FDR correction (Table S3). These GO terms reflect the presence of two primary clusters of genes. The first, which corresponds broadly to histone acetylation, may be related to chromatin remodeling and therefore is expected to effect gene expression levels across a large number of loci. Previous work focused on this ancestry cline has identified chromatin remodeling genes as a potentially important component locally adaptive variation on this ancestry cline [78]. This may indicate that this previous efforts to identify spatially varying selection in these populations may have been detecting selection on local ancestry components associated with ecological adaptation in ancestral populations. The second GO cluster, eukaryotic translation initiation factor 2 complex, also appears to implicate a central role of clinal LA outliers on the regulation of gene expression. One plausible explanation of these observations is that gene expression, particularly high level regulation of gene expression, may be especially likely to contribute to epistatic interactions as these proteins will inherently interact with a diverse set of loci throughout the genome. Given that two distinct gene clusters related to gene expression are identified by this analysis, gene expression would appear to be a plausible candidate phenotype to investigate in future work on ecological divergence and isolating factors in admixed *D. melanogaster* populations. Testing this prediction empirically through expression profiling may therefore offer fruitful grounds of understanding the earliest stages of reproductive isolation.

### Regions of Decreased African Ancestry

Another subset of loci that we may wish to identify using these data are those that contribute to reproductive isolation between African and Cosmopolitan *D. melanogaster* populations and would therefore be removed by selection from most populations on this ancestry cline. Although it is possible that Cosmopolitan alleles would be disfavored in an admixed background as well, because these populations are predominantly Cosmopolitan, we expect that the majority of selection on negatively epistatically interacting loci would remove African alleles from populations. To identify these loci, we first computed the mean African ancestry across all populations, and we then identified the subset of loci that were in the lowest 5% tail. From those loci, we selected the loci minima from adjacent genomic windows (see Methods, Figure 8), and we obtained a total of 84 local ancestry outliers.

**Figure 8.**
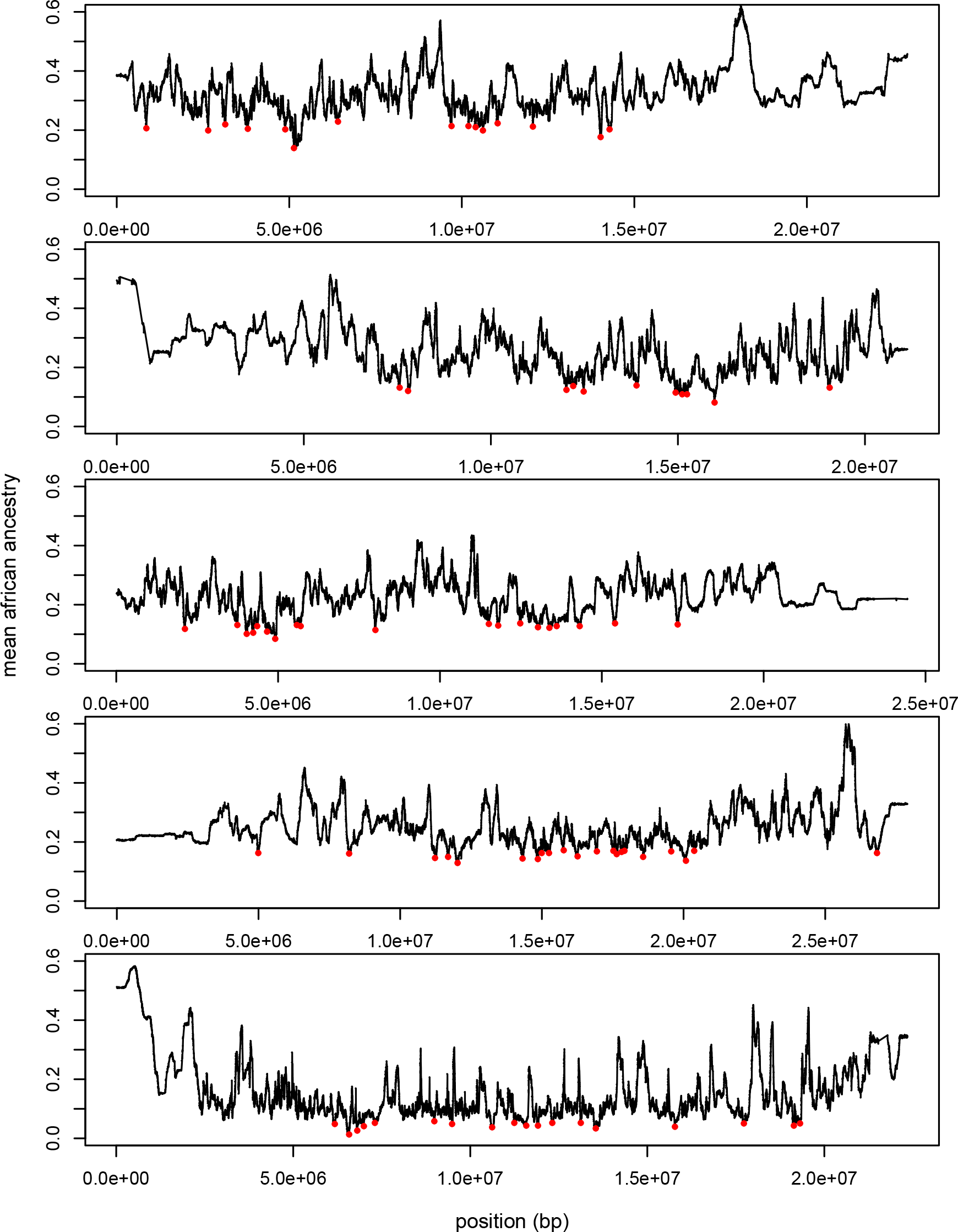
The mean African ancestry proportion across all populations on the ancestry cline for chromosome arms 2L, 2R, 3L, 3R, and X (top to bottom). Local minima outlier loci are shown in red (see Methods).

As with the clinal outlier analysis above, to identify commonalities in the types of loci identified by this analysis, we performed GO analysis on the set of loci that are outliers for the mean proportion of African ancestry. After a 5% FDR correction, there are again several gene clusters that are significantly enriched in this set of outlier loci (Supplemental Table S4). Of particular interest is the GO term oogenesis, which may indicate that female reproduction is affected during admixture between cosmopolitan and African populations of *D. melanogaster*. This finding is particularly interesting in light of the fact that female fertility is strongly affected when autosomal chromosomes from one end of this ancestry cline are made homozygous on a genetic background carrying the X chromosome from the other end of this ancestry cline [47]. Hence, the effects of combining divergence ancestry types on female fertility, and specifically the genetic basis of oogenesis, may be an appealing phenotype to characterize in detail in attempting to clarify the genetic effects that isolate African and Cosmopolitan *D. melanogaster* populations.

### Candidate Behavioral Reproductive Isolation Genes

Given the abundance of evidence supporting a role for pre-mating isolation barriers between African and Cosmopolitan flies [72–74], we are interested in highlighting genes potentially related to behavioral isolation between ancestral populations of *D. melanogaster*. Consistent with this observation, one of the strongest LA cline outliers, *egh*, has been conclusively linked to strong effects on male courtship behavior using a variety of genetic techniques [79]. Additionally, gene knockouts of CG43759, another LA cline outlier locus, have strong effects on inter-male aggressive behavior [80], and may also contribute to behavioral differences between admixed individuals. These loci are therefore appealing candidate genes for functional follow-up analyses, and illustrate the power of this LAI approach for identifying candidate genes that are potentially associated with well characterized phenotypic differences between ancestral populations.

### Little Evidence for Seasonal LA Outliers

The Pennsylvania population included in this study has been sampled extensively, including several paired fall and spring samples across three consecutive years. Previously, Bergland et al. [40] identified numerous SNPs that showed recurrent and rapid seasonal frequency changes in the Pennsylvania populations included in this study. They concluded that these sites are experiencing recurrent selection associated with recurrent environmental seasonal changes. To determine if LA across the *D. melanogaster* genome might also experience selection associated with seasonal frequency shifts, we searched for loci that showed a strong recurrent seasonal shift in LA. However, we identified fewer significantly seasonal sites than we would expect to by chance (the proportion of significant site at the alpha = 0.05 level of significance is 0.041). Furthermore, after applying a false discovery rate correction [81], there are no sites that are significantly seasonal at the q = 0.1 level. Collectively, these results indicate that LA within the Pennsylvania populations of *D. melanogaster* remains remarkably stable during seasonal environmental cycles.

Although this observation may, to a first approximation, appear to be at odds with the results reported in Bergland et al. [40], we believe that it is consistent with the model proposed in that work. Specifically, the authors suggested that long term balancing selection may maintain these seasonally favorable polymorphisms in diverse *D. melanogaster* populations and even in the ancestors of *D. melanogaster* and *D. simulants* [40]. We therefore may expect that these polymorphisms will be maintained at similar frequencies in African and Cosmopolitan populations. Hence, although the seasonal SNPs change rapidly in frequency between spring and fall [40], the LA at these sites can remain stable during seasonal fluctuations.

### Conclusion

A growing number of next-generation sequencing projects produce low coverage data that cannot be used to unambiguously assign individual genotypes, but which can be analyzed probabilistically to account for uncertainty in individual genotypes [82–84]. However, most existing LAI methods require genotype data derived from diploid individuals. Hence, there is an apparent disconnect between existing LAI approaches and the majority of ongoing sequencing efforts. In this work, we developed the first framework for applying LAI to pileup read data, rather than genotypes, and we have generalized this model to arbitrary sample ploidies. This method therefore has immediate applications to a wide variety of existing and ongoing sequencing projects, and we expect that this approach and extensions thereof will be valuable to a number of researchers. Although evaluating this application is beyond the scope of this work, one particularly enticing potential use of this method is LAI in ancient DNA samples for which sequencing depths often preclude accurate genotype calling. Importantly, it would be straightforward to model site-specific errors in this framework, which could be particularly important for ancient DNA applications [6].

For many applications, a parameter of central biological interest is the time since admixture began (*t*). A wide variety of approaches have been developed that aim to estimate *t* and related parameters in admixed populations [26,28–31,85,86]. Many of these methods are based on an inferred distribution of tract lengths, however, inference of the ancestry tract length distribution is associated with uncertainty that is typically not incorporated in currently available methods for estimating *t*. Furthermore, incorrect assumptions regarding *t* have the potential to introduce biases during LAI. Hence, it is preferable to estimate demographic parameters such as the admixture time during the LAI procedure. Nonetheless, as noted above, although LAI using our method is robust to many deviations from the assumed model, admixture time estimates are sensitive to a variety of potential confounding factors and examining the resulting ancestry tract distributions after LAI may be necessary to confirm that the assumed demographic model provides a reasonable fit to the data.

To our knowledge, this is the first method that attempts to simultaneously link LAI and population genetic parameter estimation directly, and we can envision many extensions of this approach that could expand the utility of this method to a broad variety of applications. For example, it is straightforward to accommodate additional reference populations (*e.g*. by assuming multinomial rather than binomial read sampling). Alternatively, any demographic or selective model that can be approximated as a Markov process could be incorporated—in particular, it is feasible to accommodate two-pulse admixture models and possibly models including ancestry tracts that are linked to positively selected sites. Such methods can be used to construct likelihood ratio tests of evolutionary models and for providing improved parameter estimates.

## Methods

### Constructing Emissions Probabilities

We model the ancestry using an HMM {*H*_*v*_} with state space *S* = {0,1,…,*n*}, where *H*_*v*_ = *i*, *i* ∈ *S*, indicates that in the vth position *i* chromosomes are from population 0 and *n* − *i* chromosomes are from population 1. In the following, to simplify the notation and without loss of generality, we will omit the indicator for the position in the genome as the structure of the model is the same for all positions of equivalent ploidy. We assume each variant site is biallelic, with two alleles A and a, and the availability of reference panels from source populations 0 and 1 with total allelic counts *C*_0*a*_, *C*_1*a*_, *C*_0*A*_, and *C*_1*A*_, where the two subscripts refer to population identity and allele, respectively. Also, *C*_0_ = *C*_0*A*_ + *C*_0*a*_ and *C*_1_ = *C*_1*A*_ + *C*_1*a*_. Finally, we also assume we observe a pileup of r reads from the focal population, with *r*_*A*_ and *r*_*a*_ reads for alleles A and a respectively (*r* = *r*_*A*_ + *r*_*a*_). The emission probability of state *i* ∈ *S* of the process is then defined as *e*_*i*_ = Pr(*r*_*A*_, *C*_0*A*_, *C*_1*A*_ | *r*, *C*_0_, *C*_1_, *H* = *i*, *ε*), where *ε* is an error rate. This probability can be calculated by summing over all possible genotypes in the admixed sample and over all possible population identities of the reads, as explained in the following section.

The probability of obtaining *r*_0_ (= *r* − *r*_1_) reads, in the admixed population, from chromosomes of ancestry 0, given *r* and the hidden state *H* = *i*, and assuming no mapping or sequencing biases, is binomial,

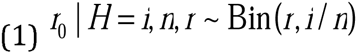

These probabilities are pre-computed in our implementation for all possible values of *i* ∈ *S* and *r*_0_, 0 ≤ *r*_0_ ≤ *r*. Similarly, for the reference populations, for *j* = 0,1,

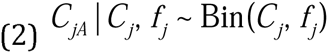

where *f*_*j*_ is the allele frequency of allele *A* in population *j*. The analogous allelic counts in the admixed population, denoted *C*_*M*0*a*_, *C*_*M*1*a*_, *C*_*M*0*A*_, and *C*_*M*1*A*_, are unobserved (only reads are observed for the admixed population), but are also conditionally binomially distributed, *i.e.*:

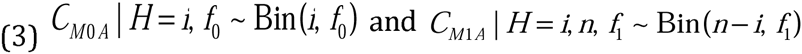

Finally, in the absence of errors, and assuming no sequencing or mapping biases, the conditional probability of obtaining *r*_*0A*_ reads of allele *A* in the admixed population is

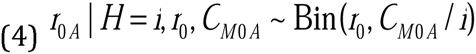

It should be noted that because we are explicitly modeling the process of sampling alleles from the population (Equation 3) and the process of sampling reads conditional on the sample allele frequencies (Equation 4), that this approach corrects for the increased variance associated with two rounds of binomial sampling in poolseq applications that has been reported previously (*e.g*., in [52]).

This probability can be expanded to include errors, in particular assuming a constant and symmetric error rate □ between major and minor allele, and assuming all reads with nucleotides that are not defined as major or minor are discarded, we have

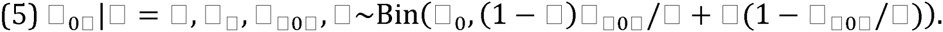

Using these expressions, and integrating over allele frequencies in the source populations, we have

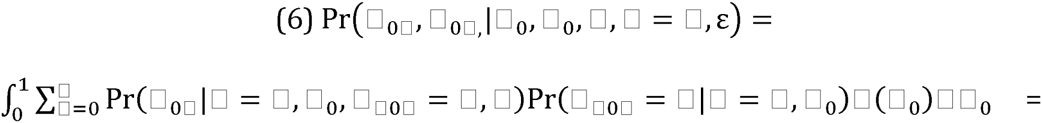

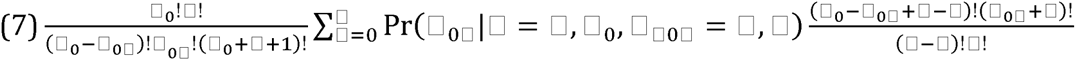

assuming a uniform [0, 1] distribution for *f*_0_. A similar expression is obtained for 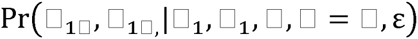 assuming *f*_1_ ~U[0,1], and these expressions combine multiplicatively to give

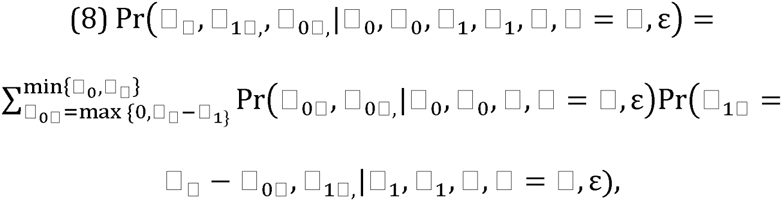

and the emission probabilities become

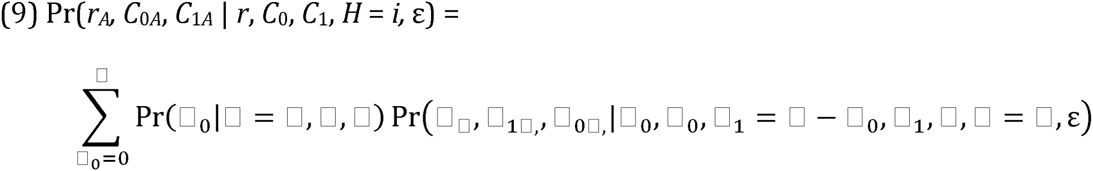

Alternatively, if the sample genotypes are known with high confidence, *i.e. C*_*MA*_ = *C*_*M*0*A*_ + *C*_*M*1*A*_ is observed, the emission probabilities are the defined as

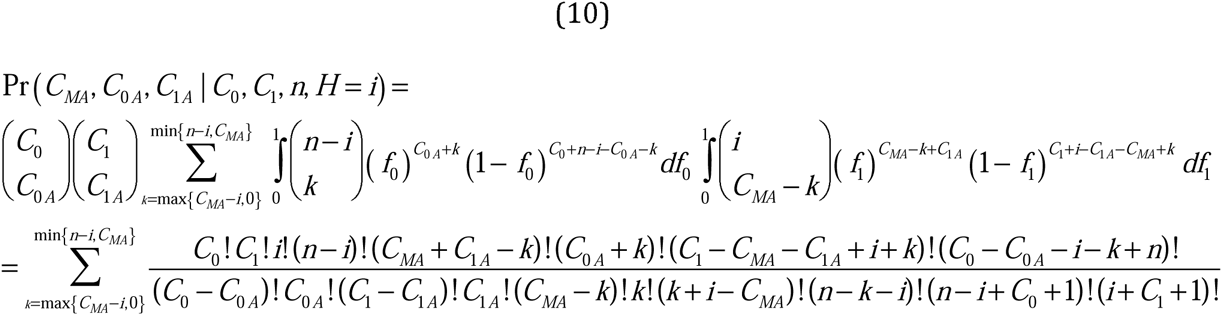

These emissions probabilities are sometimes substantially faster to compute than those for short read pileups, especially when sequencing depths are high. However, the genotypes must be estimated with high accuracy for this approach to be valid. For applications with low read coverage, or with ploidy >2 for which many standard genotype callers are not applicable, it is usually preferable to use the pileup-based approach described above.

### Constructing Transition Probabilities

We assume an admixed population, of constant size, with *N* diploid individuals, in which a proportion *m* of the individuals in the population where replaced with migrants *t* generations before the time of sampling. Given these assumptions, and an SMC’ model of the ancestral recombination graph [87], the rate of transition from ancestry 0 to 1, along the length of a single chromosome, is

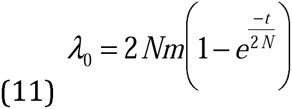

per Morgan [57]. Similarly, the rate of transition from ancestry 1 to 0 on a single chromosome is

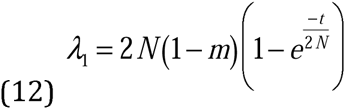

per Morgan. Importantly, because these expressions are based on a coalescence model, they account for the possibility that a recombination event occurs between two tracts of the same ancestry type and the probability that the novel marginal genealogy will back-coalesce with the previous genealogy [57]. Both events are expected to decrease the number of ancestry switches along a chromosome and ignoring their contribution will cause overestimation of the rate of change between ancestry types between adjacent markers.

The transition rates are in units per Morgan, but can be converted to rates per bp, by multiplying with the recombination rate in Morgans/bp, *r*_*bp*_ within a segment. The transition probabilities of the HMM for a single chromosome, **P**(*l*) = {*P_ij_* (*l*)},*i*, *j* ∈ *S*, between two markers with a distance l between each other, is then approximately

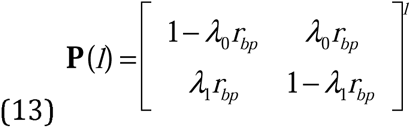

using discrete distances, or

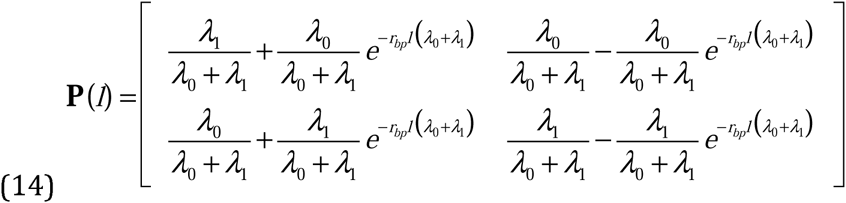

using continuous distances along the chromosome. Here, we use the continuous representation for calculations. We emphasize that the assumption of a Markovian process is known to be incorrect [57], in fact admixture tracts tend to be more spatially correlated than predicted by a Markov model, and the degree and structure of the correlation depends on the demographic model [57]. Deviations from a Markovian process may cause biases in the estimation of parameters such as *t*.

The Markov process defined above is applicable to a single chromosome. We now want to approximate a similar process for a sample of *n* chromosomes from a single sequencing pool. The true process is quite complicated, and we choose for simplicity to approximate the process for *n* chromosomes sampled from one population, as the union of *n* independent chromosomal processes. We will later quantify biases arising due to this independence assumption using simulations. Under the independence assumption, the transition probability from *i* to *j* is simply the probability of *l* transitions from state 1 to state 0 in the marginal processes and *j* − *i* + *l* transitions from state 0 to state 1, summed over all admissible values of *l*, *i.e.*,

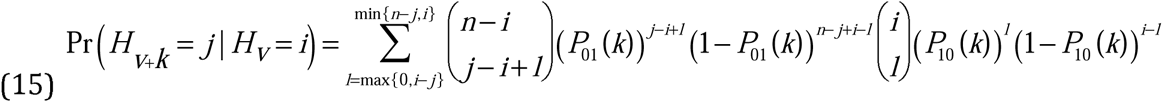

Although this procedure can be computationally expensive when there are many markers, read depths are high, and especially when *n* is large, in our implementation, we reduce the compute time by pre-calculating and storing all binomial coefficients.

### Estimating Time Since Admixture

A parameter of central biological interest, that is often unknown in practice, is the time since the initial admixture event (*t*). We therefore use the HMM representation to provide maximum likelihood estimates of *t* using the forward algorithm to calculate the likelihood function. As this is a single parameter optimization problem for a likelihood function with a single mode, optimization can be performed using a simple golden section search [88]. Default settings for this optimization in our software, including the search range maxima defaults, *t*_*max*_ and *t*_*min*_, are documented in the C++ HMM source code provided at https://github.com/russcd/Ancestry_HMM.

### Posterior Decoding

After either estimating or providing a fixed value of the time since admixture to the HMM, we obtained the posterior distribution for all variable sites considered in our analysis using the forward-backward algorithm, and we report the full posterior distribution for each marker along the chromosome.

### Simulating Ancestral Polymorphism

To validate our HMM, we generated sequence data for each of two ancestral populations using the coalescent simulator MACS [55]. We sought to generate data that could be consistent with that observed in Cosmopolitan and African populations of *D. melanogaster*, which has been studied previously in a wide variety of contexts [2,11,35–37]. We used the command line “macs 400 10000000 -i 1 -h 1000 -t 0.0376 -r 0.171 -c 5 86.5 -I 2 200 200 0 -en 0 2 0.183 -en 0.0037281 2 0.000377 -en 0.00381 2 1 -ej 0.00382 2 1 -eN 0.0145 0.2” to generate genotype data. This will produce 200 samples of ancestry 0 and 200 samples of ancestry 1 on a 10mb chromosome—*i.e.* this should resemble genotype data for about half of an autosomal chromosome arm in *D. melanogaster*. Unless otherwise stated below, we then sampled the first 50 chromosomes from each ancestral population as the ancestral population reference panel, whose genotypes are assumed to be known with low error rates. The sample size was chosen because it is close to the size of the reference panel that we obtained in our application of this approach to *D. melanogaster* (below).

To evaluate the performance of our method on data consistent with human populations, we simulated data that could be consistent with that observed for modern European and African human populations. Specifically, we simulated the model of [89] using the command line “macs 200 1e8 -I 3 100 100 0 -n 1 1.682020 -n 2 3.736830 -n 3 7.292050 -eg 0 2 116.010723 -eg 1e-12 3 160.246047 -ma x 0.881098 0.561966 0.881098 x 2.797460 0.561966 2.797460 x -ej 0.028985 3 2 -en 0.028986 2 0.287184 -ema 0.028987 3 x 7.293140 x 7.293140 x x x x x -ej 0.197963 2 1 -en 0.303501 1 1 -t 0.00069372 -r 0.00069372”. Admixture between ancestral populations was then simulated as described below.

### Simulating Admixed Populations

Although it is commonly assumed that admixture tract lengths can be modeled as independent and identically distributed exponential random variables (*e.g*. [26,29] and in this work, above), this assumption is known to be incorrect as ancestry tracts are neither exponentially distributed, independent across individuals, nor identically distributed along chromosomes [57]. We therefore aim to determine what bias violations of this assumption will have on inferences obtained from this model. Towards this, we used SELAM [56] to simulate admixed populations under the biological model described above. Because this program simulated admixture in forward time, it generates the full pedigree-based ancestral recombination graph, and is therefore a conservative test of our approach relative to the coalescent which is known to produce incorrect ancestry tract distributions for short times [57]. Briefly, we initialized each admixed population simulation with a proportion, *m*, of ancestry from ancestral population 1, and a proportion 1-*m* ancestry from ancestral population 0. Unless otherwise stated, all simulations were conducted with neutral admixture and a hermaphroditic diploid population of size 10,000.

We then assigned the additional, non-reference chromosomes from the coalescent simulations, to each ancestry tract produced in SELAM simulations according to their local ancestry along the chromosome. In this way, each chromosome is a mosaic of the two ancestral subpopulations. See, *e.g*. [2], for a related approach for simulating genotype data of admixed chromosomes.

### Pruning Ancestral Linkage Disequilibrium

Correlations induced by LD between markers within ancestral populations violates a central assumption of the Markov model framework. Although it may be feasible to explicitly model linkage within ancestral populations (*e.g*., [24,25]), when ancestral populations have relatively little LD, such as those of *D. melanogaster*, another effective approach is to discard sites that are in strong LD in the ancestral populations. Hence, to avoid this potential confounding aspect of the data, we first computed LD between all pairs of markers within each reference panel that are within 0.01 centimorgans of one another. We then discarded one of each pair of sites where |*r*| in either reference panel exceeded a particular threshold, and we decreased this threshold until we obtained an approximately unbiased estimate of the time since admixture estimates of the HMM. This approach differs from a previous method, WinPop [34], where LD is pruned from within admixed samples (see also below).

### Simulating Sequence Data

We first identified all sites where the allele frequencies of the ancestral populations differ by at least 20% within the reference panels. We excluded weakly differentiated sites to decrease runtime and because these markers carry relatively little information about the LA at a given site. Then, to generate data similar to what would be produced using Illumina sequencing platforms, we simulated allele counts for each sample, by first drawing the depth at a given site from a Poisson distribution. In most cases and unless otherwise stated, the mean of this distribution is set to be equal to the sample ploidy. We did this to ensure equivalent sequencing depth per chromosome regardless of pooling strategy, and because this depth is sufficiently low that high quality genotypes cannot be determined. We then generated set of simulated aligned bases via binomial sampling from the sample allele frequency and included a uniform error rate of 1% for both alleles at each site.

Unless otherwise stated, we simulated a total of 40 admixed chromosomes. The HMM can perform LAI on more than one sample at a time, and we therefore included all samples when running it. Hence, we used 40 haploid, 20 diploid, 4 pools of 10 chromosomes, and 2 pools of 20 chromosomes for most comparisons of accuracy reported below, unless otherwise stated. It is worth noting that it is possible to jointly analyze distinct samples from the same subpopulation that have been sequenced at different ploidies.

### Simulating Divergent Ancestral Populations

To investigate the effects of allele frequency differences between reference populations and admixed populations, we performed coalescent simulations using the software MACS [55], using the command line “./macs 500 10000000 -i 1 -h 1000 -t 0.0376 -r 0.171 -c 5 86.5 -I 8 100 100 50 50 50 50 50 50 0 -en 0 2 0.183 -en 0.0037281 2 0.000377 -en 0.00381 2 1 -ej 0.00382 2 1 -eN 0.0145 0.2 -ej 0.0005 3 2 -ej 0.000500001 4 1 -ej 0.001 5 2 -ej 0.001000001 6 1 -ej 0.002 7 2 -ej 0.002000001 8 1”. This might be expected to produce populations that are differentiated similarly to how populations of *D. melanogaster* would be across European populations or between populations in Central Africa. We then substituted the increasingly divergent populations for the reference panel. All allele frequency differences and LD pruning were performed as described above on each of the substitute reference panels.

### Accuracy Statistics

To evaluate the performance of the HMM, we computed four statistics. First, we compute the proportion of sites where the true state is within the 95% posterior credible interval, where ideally, this proportion would be equal to or greater than 0.95. As this HMM has discrete states, there are many ways the 95% credible interval could be defined. In light of the fact that the credible interval tends to be narrow (Results), we defined the interval to include all states that are overlapped, by any amount, in the 95% confidence interval of the posterior distribution. Second, we compute the mean posterior error, the average distance between the posterior distribution of hidden states and the true state

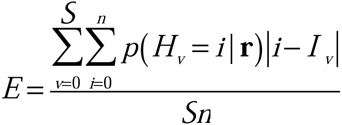

Here *S* is the total number of sites, *I_v_* is the true state at site *v*, and **r** is all the combined read data. Third, we also report the proportion of sites where the maximum likelihood estimate of the hidden state is equal to the true ancestry state. Finally, as an indicator of the specificity of our approach, we also report the average width of the 95% credible interval.

### Deviations from the Assumed Neutral Demographic Model

A potential issue with this framework is that the assumptions underlying the transition matrixes and related time of admixture estimation procedure is likely to be violated in a number of biologically relevant circumstances. We therefore simulated populations wherein individuals of ancestral population 1 began entering a population entirely composed of individuals from ancestral population 0, at a time *t* generations before the present, at a constant rate that is sustained across all subsequent generations until the time of sampling. That is, additional unadmixed individuals of ancestry 1 migrate each generation from *t* until the present.

Natural selection acting on admixed genetic regions has been inferred in a wide variety of systems (*e.g*. [5,7,13,17,18]), and is expected to have pronounced effects on the distribution of LA among individuals within admixed populations. Here again, this aspect of biologically realistic populations will tend to violate central underlying assumptions of the model assumed in this work. Towards this, we simulated admixed populations with a single pulse of admixture *t* generations prior to the time of sampling. We then incorporated selection at 2,5,10, and 20 loci at locations uniformly distributed along the length of the chromosome arm. All selected loci were assumed to be fixed within each ancestral population. Selection was additive and selective coefficients were assigned based on a uniform [0.005, 0.05] distribution to either ancestry 0 or 1 alleles with equal probability. As above, these simulations were conducted using SELAM [56].

For both selected and continuous migration simulations, we then performed the genotype and read data simulation procedure, and reran our HMM as described above. We performed 10 simulations for each treatment.

### Comparisons to WinPop

We next sought to compare our method to a commonly used local ancestry inference method, WinPop [34]. Towards this, we again simulated data using MACS and SELAM as described above. For these comparisons, the initial ancestry contribution was 0.5 and the number of generations since admixture varied between 5 and 1000. For comparison, we supplied WinPop and our program the correct time since admixture and ancestry proportions, as these are required parameters for WinPop. We also supplied the program with genotypes rather than read counts, another requirement of WinPop, whereas we supplied our HMM with read data simulated as described above. We then ran WinPop under default parameters, and we also reran WinPop using LD pruning within the reference panels, as we do in our method, instead of the default LD pruning implemented in WinPop.

### Analysis of Inuit Genotype Data

To demonstrate that LAI methods can be biased by the arbitrary selection of the time since admixture, we analyzed a dataset of SNP-array genotype data from Greenlandic Inuits. These data are described in detail elsewhere [60,61]. This population has received some admixture from a European source population, and the authors had previously used RFMix [24] to perform LAI, and found some sensitivity to the assumed time since admixture (J. Crawford *pers. Comm.*). We analyzed data from chromosome 10 using RFMix v1.5.4 [24] as described in Moltke *et al.* [61] assuming admixture occurred either 5 or 20 generations ago. We subsequently analyzed chromosome 10 using our HMM including the genotype-analysis emissions probabilities and assuming a genotype error rate of 0.2%. For our analysis we identified the LD cutoff that is appropriate for these data as described above.

### Generating *D. melanogaster* Reference Populations

To generate reference panels, we used a subset of the high quality D. melenaogaster assemblies that have been described previously in Pool *et al*. (2012) and Lack *et al*. (2015). As in the local ancestry analysis of Pool (2015), we used the French population. For our African reference panel, we selected a subset of the Eastern and Western African populations (CO, RG, RC, NG, UG, GA, GU) and grouped them into a single population for the purposes of our analysis. We elected to combine populations so that we would have a larger reference panel of African populations for this analysis, this solution may be justified because these *D. melanogaster* populations are only weakly genetically differentiated [2,21,90], particularly after common inversion-bearing chromosomes are removed from analyses. Specific individuals were selected for inclusion in the African reference panel if previous work found they have relatively little cosmopolitan ancestry (*i.e.*, below 0.2 genome-wide in [2]).

Because of their powerful effects on recombination, chromosomal inversions are known to have substantial impacts on the distribution of genetic variants on chromosomes containing chromosomal inversions in *D. melanogaster* [2,63]. For this reason, we removed all common inversion-bearing chromosome arms from the reference populations [91]. Nonetheless, it is clear that chromosomal inversions are present in the pool-seq samples [64]. Although the inversions certainly violate key assumptions of our model—particularly the transmission probabilities—given that our approach is robust to a many perturbations, we expect the LA within inverted haplotypes can be estimated with reasonable confidence, and the overall LAI procedure will still perform adequately with low frequencies of chromosomal-inversion bearing chromosomes present within these samples.

Although these reference populations are believed to have relatively little admixture, some admixture is likely to remain within these samples [2]. To mitigate this potential issue, we first applied our HMM to each reference population using the genotype-based emissions probabilities (above). Calculated across all individuals, we found that our maximum likelihood ancestry estimates were identical with those of Pool *et al*. (2012) at 96.2% of markers considered in our analysis. The differences between the results of these methods may reflect differences in the methodology of LAI or differences in the reference panels. Nonetheless, the broad concordance suggests the two methods are yielding similar overall results. We masked all sites where the posterior probability of non-native ancestry was greater than 0.5 within each reference individual’s genome. These masked sequences were then used as the reference panel for the analyses of pool-seq data below.

### Ancestry Cline Sequence Data Analysis

We acquired pooled sequencing data from six populations from the east coast of the United States. The generation of these samples, sequencing data, and accession numbers are described in detail in [21,40]. Briefly, the samples are comprised of individuals drawn from natural populations and sequenced in relatively large pools of 66–232 chromosomes. We aligned all data using BWA v0.7.9a-r786 [92] using the ’MEM’ function and the default program parameters. For all alignments, we used version 5 of the *D. melanogaster* reference genome [93] in order to make our analysis and coordinates compatible with the *Drosophila* genome nexus [91]. We then realigned all reads using the indelrealigner tool within the GATK package [84], and we extracted the sequence pileup using samtools mpileup v1.1 [94] using the program’s default parameters.

We extracted sites at ancestry informative positions within the reference panels, where we required that the reference panel have a minimum of 50% of individuals with a high quality genotype call in both Cosmopolitan and African reference populations. As above, ancestry informative sites were defined as those with a minimum of 20% difference in allele frequencies between the reference panels used, and we retained only ancestry informative sites for our analyses. We then produced global ancestry estimates for each chromosome arm separately for each sample using the method of Bergland *et al.* (2016). We ran our HMM for each chromosome arm and each population, and we provided the program this estimate of the ancestry proportion and the time since admixture, 1593 generations [17]. We elected to provide the time since admixture because we have found that this parameter is difficult to estimate in relatively large pools (see Results). However, the program can accurately estimate LA in high ploidy samples even when the time since admixture cannot be estimated correctly (see Results).

### Correlation with Local Recombination Rates

To assess the correlation between local recombination rates and LA in the genome, we computed Spearman’s rank sum correlation between the proportions African ancestry and the local recombination rates in windows of 100 ancestry informative markers. As above, we used the recombination rate estimates of [59]. We estimated confidence intervals using 1000 block-bootstrap samples using window sizes of 100 SNPs.

### Robustness of LAI to Genomic Heterogeneity

To determine if there are systematic biases in LAI across the genome, we computed the mean difference in genomic windows between LA estimates for two samples form Maine and between two samples from Florida. We assessed evidence for systematic biases through the correlation between local recombination rates and differences in local ancestry inference using Spearman’s rank sum correlation.

### Identifying LA Cline Outliers

To detect loci that show evidence for steeper ancestry clines than the genomic average, we first computed the Spearman’s rank correlation between mean ancestry proportions and latitude for each chromosome arm separately. Then, for each site for which we obtained a posterior ancestry distribution for all samples, we computed the partial Spearman’s rank correlation between the posterior ancestry mean and latitude while correcting for the correlation between latitude and the overall ancestry proportion. We then computed the probability of obtaining the observed partial correlation in R, which implements the approach of [95], and we retained those sites where the probability of the partial correlation between local ancestry and latitude was less than 0.005 as significant in our analysis. Although this cutoff is arbitrary, given the strong evidence for local adaptation and reproductive isolation in these populations [46,47,96], the tail of the LA cline distribution will likely be enriched for sites experiencing selection on this ancestry gradient. Due to linkage, adjacent sites show strong autocorrelation. We therefore selected the local optima for a given clinally significant LA segment (*i.e.* a tract where all positions are significantly correlated with latitude at our threshold) and retained these for analyses of outlier loci. Finally, to further reduce the effect of autocorrelation, we retained only those local optima for which no other optimum had a stronger correlation with latitude within 100,000bp on either side on the site.

### Identifying Low African Ancestry Outlier Loci

To identify loci with a disproportionately low proportion of African ancestry across this ancestry cline, we computed the mean African ancestry across all populations. We then selected those sites in the lowest 5% tail on each chromosome arm and selected only the local minima within 100kb windows on either side of a selected locus.

### Gene Ontology Analyses

We performed Gene-ontology (GO) analyses on outlier SNPs using Gowinda [97], where the background set of SNPs was all positions at which we obtained a posterior distribution in all samples (*i.e.* the set on which we obtained estimates of the posterior probability of African ancestry). We ran the program using default parameters, except that we included all genes within 10000bp of a focal SNP, and we performed 1e6 total GO simulations.

### Seasonality of LA in the Pennsylvania Populations

To identify recurrent seasonal changes in the local ancestry, we followed an approach similar to [40]. Specifically, we fit a generalized linear model of the form

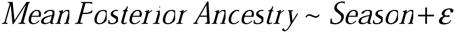

We then recorded the estimated effect size, and probability of the observed correlation for each site in the genome at which we obtained a posterior ancestry distribution in all samples considered. To correct for multiple testing, we applied a false discovery rate correction [81] to the resulting p-value distribution.

## Acknowledgements

We thank three anonymous reviewers, Tyler Linderoth, Shelbi Russell and members of the Nielsen and Slatkin labs for helpful comments. We also thank Jacob Crawford for providing Inuit polymorphism data and for providing the observation that LAI inference depended heavily on *t*.

---

**Supplemental Figure S1.** Comparison between LAI using the full ancestral recombination graph via forward-time simulations (red) with those from independent and identically distributed draws from the SMC’ distribution (black). Simulations were conducted using an ancestry proportion of 0.25 and population size of 10,000 hermaphroditic individuals.

**Supplemental Figure S2.** Effects of unknown admixed population sizes on LAI. All LAI was conducted assuming the true population size was 10,000. Simulated population sizes were 100 (black), 1,000 (red), 10,000 (blue) and 100,000 (green). Ploidy 1 on the right, ploidy 2 on the left. From top to bottom, rows are the estimated time of admixture, the proportion of sites where the true state is within the 95% credible interval, the width of the 95% credible interval, the mean posterior error, and the proportion of times that the maximum likelihood estimate is equal to the true state. For all simulations, the ancestry proportion was equal to 0.5.

**Supplemental Figure S3.** LAI accuracy when admixture times are increasingly ancient. Here, ancestry proportions are 0.5 (black), 0.25 (blue), 0.1 (violet), 0.75 (orange) and 0.9 (red). From top to bottom, statistics plotted are estimated time, the proportion of sites where the true ancestry frequency is within the 95% credible interval, the mean 95% credible interval width, mean posterior error, and the proportion of times that the maximum likelihood estimate is correct.

**Supplemental Figure S4.** The effects of reference panel size on LAI and time estimation using the HMM. Here, we compare reference panels of size 100 (blue) with reference panels of size 10 (black). From left to right, ancestry proportions are 0.1, 0.25, 0.5, 0.75 and 0.9. From top to bottom the plotted statistics are estimated time, proportion in the 95% credible interval, the average width of the 95% credible interval, the mean posterior error, and the proportion of sites where the maximum likelihood ancestry estimate is correct.

**Supplemental Figure S5.** Accuracy of time estimation and LAI when reference populations are increasingly divergent from the source of the admixture pulses. In columns are divergence times between ancestral populations (in units of 4Ne) of 0, 0.0005, 0.001, 0.002. From top to bottom the plotted statistics are estimated time, proportion in the 95% credible interval, the average width of the 95% credible interval, the mean posterior error, and the proportion of sites where the maximum likelihood ancestry estimate is correct.

**Supplemental Figure S6.** Comparison of the proportion of sites where the maximum likelihood ancestry estimate of local ancestry is correct between WinPop and our method. WinPop was run with default parameters (black), and with LD pruned in the ancestral populations, but not in the admixed population (red). Our method was run with default parameters (blue), but with the time since admixture and correct ancestry proportion supplied to our program as these parameters are required by WinPop.

**Supplemental Figure S7.** Bias in LAI due to uncertainty in *t*. The posterior probability estimated using RFMix of European ancestry at a given site in the genome assuming t = 5 (black) and assuming t = 20 (red) for a sample representative of the average difference (top left) and a more extreme example (top right). The distribution of differences in mean Inuit ancestry for all samples (bottom left) using RFMix. The log likelihood of each time since admixture as computed using our method (bottom right), which shows a clear optimum at 6-7 generations since admixture. All analyses were restricted to SNPs on chromosome 10.

**Supplemental Figure S8.** Bias in LAI and time estimation due to incorrect estimation of *m*. On the left, true m is 0.1and on the right true m is 0.5. Supplied m varies across 0.05 to 0.95. From top to bottom, the plotted statistics are estimated t, proportion in the 95% confidence interval, mean 95% confidence interval width, mean posterior error and the proportion of sites where the maximum likelihood estimate is correct. All plots include ploidy one (back), ploidy two (red), ploidy ten (blue), and ploidy twenty (green).

**Supplemental Figure S9.** Bias in LAI and time estimation due to incorrect assumptions of *t*. On the left, true *t* is 100 and on the right true *t* is 1000. Supplied *t* varies across 100 to 2000 generations. From top to bottom, the plotted statistics are estimated t, proportion in the 95% confidence interval, mean 95% confidence interval width, mean posterior error and the proportion of sites where the maximum likelihood estimate is correct. All plots include ploidy one (back), ploidy two (red), ploidy ten (blue), and ploidy twenty (green).

**Supplemental Figure S10.** Estimates of *t* obtained from block bootstrap replicates for populations that have admixed for 1000 (top), and 2000 (bottom) generations. From left to right, sample ploidies are 1, 2, 10, and 20. For both simulations, *m* = 0.5.

**Supplemental Table S1.** Comparison of run times for various demographic models and sample ploidies using this method.

**Supplemental Table S2.** LA clinality in the genomic intervals immediately surrounding breakpoints of known polymorphic inversions.

**Supplemental Table S3.** Results of GO analysis of 80 identified LA clinal outlier loci.

**Supplemental Table S4.** Results of GO analysis of 84 identified low African ancestry outlier loci.

